# IGF2BP1 induces high-risk neuroblastoma and forms a druggable feedforward loop with MYCN promoting 17q oncogene expression

**DOI:** 10.1101/2023.03.18.533148

**Authors:** Sven Hagemann, Danny Misiak, Jessica L. Bell, Tommy Fuchs, Marcell I. Lederer, Nadine Bley, Monika Hämmerle, Ehab Ghazy, Wolfgang Sippl, Johannes H. Schulte, Stefan Hüttelmaier

## Abstract

**Background:** Neuroblastoma is the most common solid tumor in infants accounting for approximately 15% of all cancer-related deaths. Over 50% of high-risk neuroblastoma relapse, emphasizing the need of novel drug targets and therapeutic strategies. In neuroblastoma, chromosomal gains at chromosome 17q, including *IGF2BP1*, and *MYCN* amplification at chromosome 2p are associated with adverse outcome. Recent, pre-clinical evidence indicates the feasibility of direct and indirect targeting of IGF2BP1 and MYCN in cancer treatment.

**Methods:** Candidate oncogenes on 17q were identified by profiling the transcriptomic/genomic landscape of 100 human neuroblastoma samples and public gene essentiality data. Molecular mechanisms and gene expression profiles underlying the oncogenic and therapeutic target potential of the 17q oncogene *IGF2BP1* and its cross-talk with *MYCN* were characterized and validated in human neuroblastoma cells, xenografts and PDX as well as novel IGF2BP1/MYCN transgene mouse models.

**Results:** We reveal a novel, druggable feedforward loop of IGF2BP1 (17q) and MYCN (2p) in high-risk neuroblastoma. This promotes 2p/17q chromosomal gains and unleashes an oncogene storm resulting in fostered expression of 17q oncogenes like BIRC5 (survivin). Conditional, sympatho- adrenal transgene expression of IGF2BP1 induces neuroblastoma at a 100% incidence. IGF2BP1- driven malignancies are reminiscent to human high-risk neuroblastoma, including 2p/17q-syntenic chromosomal gains and upregulation of Mycn, Birc5, as well as key neuroblastoma circuit factors like Phox2b. Co-expression of IGF2BP1/MYCN reduces disease latency and survival probability by fostering oncogene expression. Combined inhibition of IGF2BP1 by BTYNB, MYCN by BRD inhibitors or BIRC5 by YM-155 is beneficial *in vitro* and, for BTYNB, also *in vivo*.

**Conclusion:** We reveal a novel, druggable neuroblastoma oncogene circuit settling on strong, transcriptional/post-transcriptional synergy of MYCN and IGF2BP1. MYCN/IGF2BP1 feed-forward regulation promotes an oncogene storm harboring high therapeutic potential for combined, targeted inhibition of IGF2BP1, MYCN expression and MYCN/IGF2BP1-effectors like BIRC5.

## Background

Neuroblastoma is the most common extracranial childhood tumor and originates from precursors of the sympatho-adrenal cell lineage [1]. High-risk neuroblastoma (HRN) accounts for approximately 15% of all cancer-related death in infants and despite intensive multimodal therapy >50% of HRN relapse [2]. This emphasizes the need to elucidate novel drug targets and therapeutic avenues. Neuroblastoma is commonly associated with genomic abnormalities including *MYCN* amplification (∼25-35%; Chr 2p), 1p and 11q loss as well as 17q gain, the most frequent (>50%) chromosomal aberration [1–4]. Intriguingly, 17q gain is frequently associated with *MYCN* amplification and adverse disease outcome, but genetic synergies of 2p and 17q remain largely unknown [4]. Candidate oncogenes at 17q are suggested by the fact that gains at 17q are recurrent events also in other cancers, e.g. breast cancer [5] or medulloblastoma [6], and high passage human embryonic stem cells (hESCs) [7]. Two main issues have restricted the use of 17q gain in cancer management, a robust cut- off criterion for gains and only few confirmed and druggable 17q target genes. However, several studies identified 17q oncogene candidates, including *TBX2*, *PPM1D*, *TOP2A*, *BIRC5* and *ALYREF* [8–13]. Another 17q gained candidate gene, *IGF2BP1*, was reported to associate with poor neuroblastoma outcome [14, 15]. The insulin-like growth factor 2 mRNA-binding protein 1 (IGF2BP1) is an oncofetal RNA-binding protein with reported gene gain and suggested MYCN-associated roles in neuroblastoma [14]. The main function of IGF2BP1 in cancer is the partially m^6^A- (N^6^- methyladenosine), but typically microRNA- (miRNA) and 3’UTR-dependent stabilization of mRNAs [16–21]. This results in elevated expression of oncogenic factors like LIN28B, MYC and E2F1 [22–24]. IGF2BP1 is upregulated in higher-risk clinical neuroblastoma groupings, including *MYCN*-amplified (MNA) or INSS-4 (International Neuroblastoma Staging System) tumors [14]. IGF2BP1 is *de novo* expressed in several cancers and shows conserved association with poor prognosis, disease progression and metastasis [21]. However, it remains unknown if IGF2BP1 is an oncogene in human cancers, how it synergizes with MYCN in neuroblastoma, and if the targeting of IGF2BP1 harbors therapeutic benefits in cancer treatment.

Here, we reveal that IGF2BP1 is a *bona fide* druggable oncogene in neuroblastoma. For the first time, we demonstrate, that sympatho-adrenal transgene expression of IGF2BP1 is sufficient to induce HRN characterized by 2p/17q-syntenic chromosomal gains and upregulation of Mycn. MYCN and IGF2BP1 synergize by transcriptional/post-transcriptional feedforward regulation, which unleashes an oncogene storm, including upregulation of 17q genes like BIRC5/Birc5. In neuroblastoma cell and xenograft models, IGF2BP1 deletion and inhibition by the small molecule BTYNB [25] impair tumor growth. BTYNB robustly disrupts MYCN/IGF2BP1 synergy, downregulates 17q oncogenes and is beneficial with both, BIRC5 inhibition via YM-155 [26] and MYCN inhibition by the BRD-inhibitor Mivebresib [27].

## Methods

### RNA-seq library preparation and sequencing

Library preparation of human neuroblastoma tumor samples was performed on fragmented RNA. A pool of up to 10 libraries was used for cluster generation at a concentration of 10 nM using an Illumina cBot. High-throughput sequencing of 100 bp long unstranded paired-end reads was performed with an Illumina HiScanSQ sequencer at the sequencing core facility of the IZKF (Leipzig) using version 3 chemistry and flowcell according to the instructions of the manufacturer. For RNA-seq library preparation of transgenic mouse tumor samples, 2 µg of total RNA served as input for polyA(+)-RNA enriched and strand-specific library preparation, performed by Novogene (Hong Kong). Sequencing was accomplished with an Illumina NovaSeq 6000 machine.

### Shallow whole genome sequencing

A total amount of 50 ng - 1 μg DNA per sample was used as input material for DNA sample preparation of human neuroblastoma and transgenic mouse tumor samples. Shallow WGS (sWGS) of all samples was performed on an Illumina NovaSeq 6000 at Novogene (Hong Kong), resulting in a low coverage of around 0.7x per sample.

Analysis of high-throughput sequencing data

The processed sequencing reads were aligned by HiSat2 (version 2.1.0) [28] to the human (UCSC hg38) or mouse genome (UCSC mm39). SmallRNA-seq reads were aligned by Bowtie2 (version 2.3.5.1) [29] to the mouse genome (UCSC mm10). RNA-seq reads of transgenic mouse tumor samples was corrected for a determined batch effect using ComBat-seq from R package sva (version 3.46.0) [30]. Differential gene expression was determined using the R package edgeR (version human: 3.32.0, mouse: 3.40.0) [31] utilizing trimmed mean of M- values (TMM) [32] normalization. A false discovery rate (FDR) value below 0.05 was considered as threshold for the determination of differential gene expression. Principal component analysis (PCA) on normalized fragments per kilobase of transcript per million fragments mapped (FPKM) and filtered genes with zero expression was performed by the prcomp function within the R environment. R package factoextra (https://rpkgs.datanovia.com/factoextra/; version 1.0.7) was used for visualization of PCA results. Copy number (CN) variations from sWGS data was determined using the R package cn.Mops (version human: 1.36.0, mouse: 1.44.0) [33]. For determination of the ADRN/MES signature for mouse tissues, previously published gene sets were used [33]. Human ADRN/MES gene symbols were homology converted to mouse gene symbols by using the R package biomaRt (version 2.54; Supplementary Table 1) [34]. Public neuroblastoma sequencing data were analysed through the web application provided in the respective publication (https://www.neuroblastomacellatlas.org/) [35].

### Quantitative analyses of copy number data

For copy number analysis, threshold values were set to known log2 ratios of -0.4 and +0.3 associated with losses and gains, respectively [36]. Identification of balanced and unbalanced 17q samples is based on median CN fold change comparing the whole 17p region (17p) with the 17q region from IGF2BP1 locus to 17q-ter (IMP1-ter). A balance value (bv) is determined on linear regression model, separating each sample into balanced or unbalanced 17q. Linear regression is defined as bv = (-IMP1-ter + 17p + 0.2), resulting in unbalanced if bv ≤ 0 and balanced if bv > 0. Identification of *MYCN*-amplified samples is based on CN of the *MYCN* locus (CN > 4 as MNA). Visualization of CN changes and comparison of gain and loss regions was performed with the R package karyoploteR (version 1.24.0) [37]. Genome conversion of coordinates and annotations between mouse and human assemblies was performed using USCS Lift Genome Annotations (https://genome.ucsc.edu/cgi-bin/hgLiftOver).

### Pan-cancer loss-of-function analysis

To identify essential genes in MNA neuroblastoma, pan-cancer loss-of-function CRISPR screens of 620 cancer cell lines from different primary diseases were utilized for dependency analysis, using the Broad Institute Cancer Dependency Map (DepMap) portal (version 19Q3) [38]. 13 MNA cell lines (MYCN CN > 4) were selected from 46 available neuroblastoma cell lines and compared against 607 non-neuroblastoma cell lines (referred to as others). Subsequently, all available genes were filter by minimum expression (log2 TPM > 2) in at least 11 of 13 MNA cell lines (∼ 85%). Median dependency was calculated for remaining genes across MNA and others, filtering for essential genes in both groups by a median below -0.2 and above -0.3 for MNA and others, respectively. Following, genes not present in both groups or to be considered as common essential genes, retrieved by DepMap, were removed. For the final identification of MNA essential genes in neuroblastoma, significance in dependency changes between MNA and other cell lines were determined, performing a two-group comparison across the remaining genes by parametric empirical Bayes method provided by the Limma R package [39]. A FDR value below 0.05 was considered as threshold for significant essential genes in MNA (Supplementary Table 2).

### Survival analysis

Survival analysis were performed using TMM normalized expression data. MRNAs were expression filtered (at least 1 FPKM in sum across all samples) and separated by the respective median of log2-transformed expression values. Log-rank test was performed with R package survival (version 3.2-11, https://CRAN.R-project.org/package=survival). Survival analysis for IGF2BP1 and MYCN expression was determined by best cut-off method. Multi-variate cox regression was performed using the coxph funtion of the R package survival (version 3.4.0) via an in-house R script. Survival analysis upon *MYCN* amplification and/or 17q unbalancing was performed based on determined CN. Of 100 tumors one was lacking survival data and for three samples RNA sequencing failed, resulting in different amount of total samples depending on analysis (Fig. 1; Supplementary Fig. 1, 8).

**Fig. 1:**
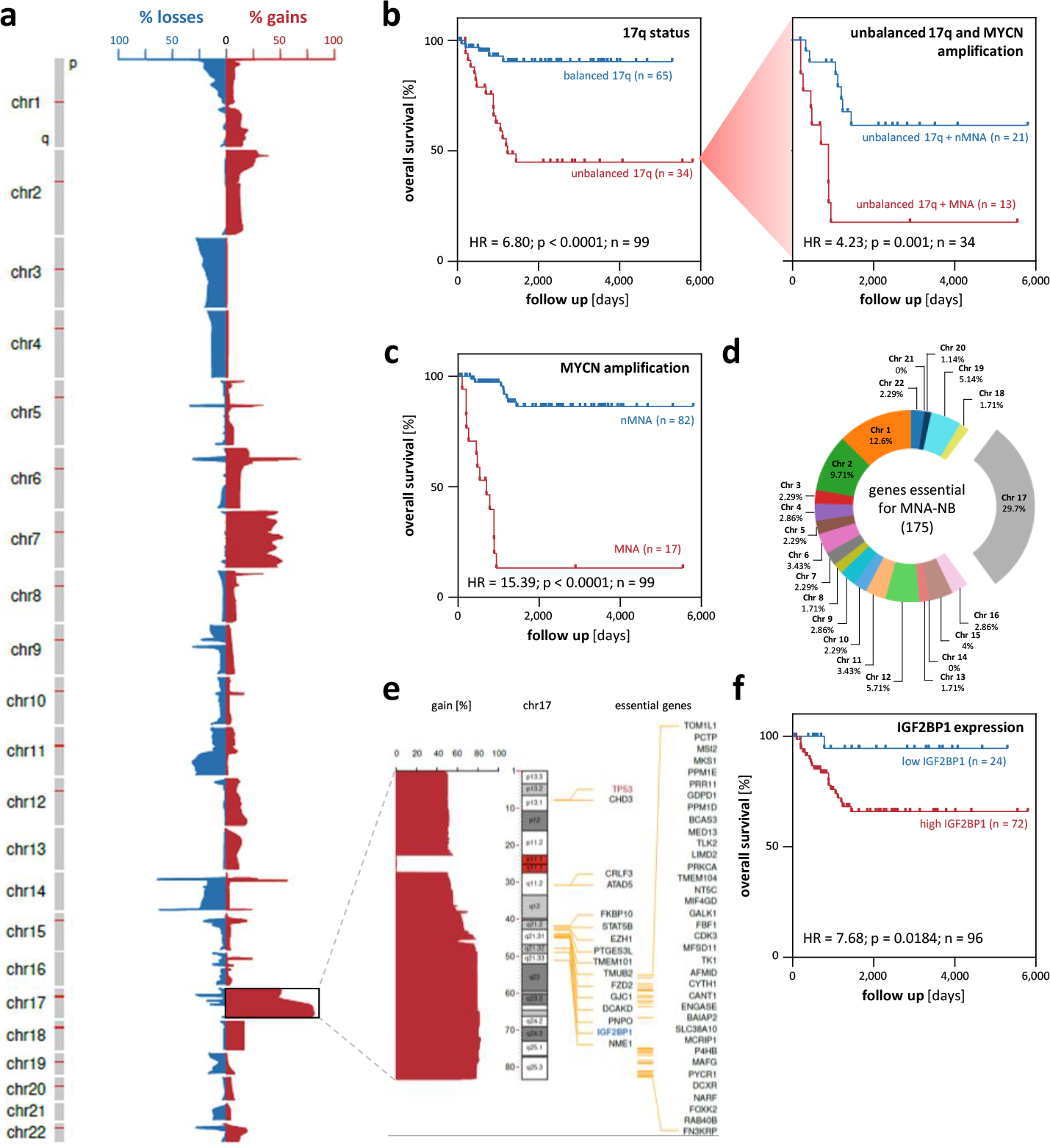
Unbalanced Chromosome 17q upregulates MNA neuroblastoma essential genes and indicates adverse disease outcome. **(a)** Frequency (%) of DNA copy number gains (red) and losses (blue) for chromosome 1 to 22 in 100 primary human neuroblastoma samples. (b, c) Kaplan-Meier survival analyses by Chr 17q balance status (b, left), MYCN amplification status in Chr 17q unbalanced tumors (b, right) and MYCN amplification status **(c)**. **(d)** Genomic distribution of 175 autosomal essential genes in MNA neuroblastoma. **(e)** Genomic location and percentage gain of essential (black and IGF2BP1 in blue) neuroblastoma genes on Chr 17 in primary neuroblastoma (non-essential TP53 on 17p is highlighted red). **(f)** Kaplan-Meier survival analyses by IGF2BP1 mRNA expression (best cut- off).

Gene set enrichment analysis. Gene set enrichment analysis (GSEA) [40] was performed using the R package clusterProfiler (version 4.6.0) [41] and MSigDB (version 2022.1) [42] gene sets with a minimum set size of 10 and no upper size restriction (Supplementary Table 3). Mouse gene symbols were homology converted to human symbols by using the R package biomaRt (version 2.54) [34].

### IGF2BP1 CLIP studies

IGF2BP1 eCLIP (enhanced crosslinking and immunoprecipitation; Supplementary Table 4) peak data of Hep-G2 and K-562 cells were obtained from the ENCODE portal (www.encodeproject.org; ENCODE Project Consortium, 2012; identifiers ENCFF486BXN, ENCFF976DBP and ENCFF435MEM, ENCFF701YCW, respectively). IGF2BP1 eCLIP data of hESCs were obtained from the Gene Expression Omnibus (GEO; sample IDs GSM2071742 and GSM2071745).

### RNA modification analysis

N^6^-Methyladenosine (m^6^A) modification sites were identified using the RNA Modification Database, RMBase (version 2.0) [43].

### MYCN ChIP-seq analysis

MYCN ChIP-seq (chromatin immunoprecipitation sequencing; Supplementary Table 5) data was obtained from ChIP-Atlas [44] database for 9 untreated neuroblastoma-derived cell lines (BE(2)-C, CHP-134, COG-N-415, KELLY, LA-N-5, NB-1643, NB69, NGP, SK-N-BE(2)), comprising 15 experiments (SRX1690205, SRX5662024, SRX5662025, SRX5662026, SRX6935370, SRX1690210, SRX2550934, SRX3542258, SRX6935374, SRX2550935, SRX6935376, SRX6935379, SRX1690213, SRX2550933, SRX1178181).

### E-Box motif locations

Locations of E-Box sequences were determined via an in-house R script using pattern matching of the MYC/N E-Box motif (CANNTG) at investigated sequences [45].

IGF2BP1 and transcription factor correlation analysis. For distinct transcription factors and IGF2BP1 the Spearman correlation coefficients with all annotated genes were determined in two independent human neuroblastoma datasets (the here presented and Kocak dataset from R2). Correlation coefficients for IGF2BP1 were then plotted against the correlation coefficients for a specific transcription factor to analyze the correlation between the IGF2BP1- and transcription factor-associated gene expression signature.

### Cell culture and siRNA transfection

BE(2)-C (ATCC, CRL-2268) cells were cultured in a 1:1 mixture of DMEM/F12 (with HEPES, Gibco) and EMEM (ATCC); KELLY (DSMZ, ACC 355), SHEP-TET21N (legal MTA agreement from Prof. Schwab at the DKFZ, CVCL_9812) and HEK293T17 (ATCC, CRL-11268) cells were cultured in DMEM (Gibco) and NBL-S (DSMZ, ACC-656) cells were cultured in IMDM (Gibco) all supplemented with 10% FBS. Cells were grown at 37°C and 5% CO_2_. Neuroblastoma cell lines (BE(2)- C, KELLY, SHEP-TET21N, NBL-S) were authenticated at Eurofins Genomics by 16 independent PCR systems (D8S1179, D21S11, D7S820, CSF1PO, D3S1358, TH01, D13S317, D16S539, D2S1338, AMEL, D5S818, FGA, D19S433, vWA, TPOX and D18S51) and compared to online databases of the DSMZ and Cellosaurus. For siRNA knockdown 4 x 10^5^ BE(2)-C, 1.5 x 10^6^ KELLY or 5 x 10^5^ TET21N cells were transfected on a 6-well plate using 7.5 µl Lipofectamine RNAiMAX (Thermo Fisher Scientific) and 50 pmol siRNA according to the manufactureŕs instructions. For the production of lentiviral particles, 3.6 x 10^6^ HEK293T17 cells were transfected on a 6-well plate with 7 µl Lipofectamine 3000 (Thermo Fisher Scientific), the packaging plasmids psPax2 (3 µg) and pMD2.G (1.5 µg) as well as the lentiviral expression vector pLVX (4 µg) encoding iRFP. 2 µl P3000 per µg DNA were added. For luciferase reporter studies 5 x 10^4^ BE(2)-C or 1.25 x 10^5^ KELLY or NBL-S cells were transfected on a 24-well plate with 1 µl Lipofectamine 2000 (Thermo Fisher Scientific) and 200 ng pmirGLO vector. For genomic deletion of IGF2BP1 via CRISPR/Cas9 5 x 10^5^ cells were transfected on a 6-well plate using 3.75 µl Lipofectamine 3000, 2 µg Cas9- and 1 µg sgRNA-encoding plasmids. For mRNA or protein decay analyses cells were treated with actinomycin D (5 µM, Sigma Aldrich) or emetin (100 µM, Sigma Aldrich) respectively for indicated time points. Plasmids and siRNAs used are summarized in Supplementary Table 6.

CRISPR/Cas9 knockout generation. For the CRISPR/Cas9-mediated genomic deletion of IGF2BP1, cells were transfected with two CRISPR sgRNA-encoding (psg-RFP-IGF2BP1 Exon6 and psg-RFP- IGF2BP1 Exon7) and a Cas9 nuclease-encoding (pcDNA-Cas9-T2A-GFP) plasmid. 48h post transfection cells were sorted for single cell clones by seeding one RFP- and GFP-positive cell per 96-well using a FACS Melody sorter (BD Bioscience). The deletion of IGF2BP1 was validated by western blotting. CRISPR sgRNAs and plasmids are summarized in Supplementary Table 6.

### Lentiviral transduction

HEK293T17 cells were transfected as described in *Cell culture and siRNA transfection*. Lentiviral particle-containing supernatant was collected 24 and 48 h post transfection and stored at -80 °C. Titers were analyzed 72 h post infection of 5 x 10^4^ HEK293T17 cells and determined by flow cytometry (iRFP) using a MACS Quant Analyzer (Miltenyi BioTech). Lentiviral transduction for downstream experiments was accomplished at 5 MOI (multiplicity of infection).

### Luciferase assay

The MYCN and BIRC5 3’UTR was amplified from genomic DNA and cloned in the pmirGLO plasmid. Mutated MYCN 3’UTR was designed by mutation of following miRNA seed regions: miR-34a, miR-19, miR-29, miR-101, let-7 and miR-17-92 and ordered from GenScript Biotech. 36 h post transfection, Dual-Glo luciferase reporter analyses were performed according to manufactureŕs protocol. Ratios of Firefly to Renilla activity were calculated and the activity of the 3’UTR reporters were normalized to the respective controls. For luciferase assays with BTYNB, medium was changed 6 h post transfection and DMSO or BTYNB was added at EC_50_ concentration. Reporter containing a minimal vector-encoded 3’UTR served as normalization control. Primers and plasmids are summarized in Supplementary Table 6.

### Spheroid formation and anoikis assay

For spheroid growth assay 2 x 10^3^ cells were seeded in a 96- well round-bottom ultra-low attachment plate (Corning) with 10% FBS. Spheroid formation was induced by centrifugation for 5 minutes at 2000 rpm. Five days post seeding images were acquired using an IncuCyte S3 (Sartorius). For anoikis assay 1 x 10^3^ cells were seeded in a 96-well flat-bottom ultra-low attachment plate (Corning) with 1% FBS. Five days post seeding cells were transferred to a round-bottom plate, centrifuged for 5 minutes at 2000 rpm and then images were acquired using an IncuCyte S3 (Sartorius). Cell viability and caspase 3/7 activity was measured with CellTiter Glo (Promega) or Caspase Glo (Promega) respectively according to manufactureŕs protocol.

### RNA immunoprecipitation

For RNA immunoprecipitations (RIP) cell extracts (3.5 x 10^6^ BE(2)-C or 12.5 x 10^6^ KELLY and NBL-S) were prepared on ice using RIP buffer (10 mM HEPES, 150 mM KCl, 5 mM MgCl_2_, 0.5% NP40, pH 7.0). Cleared lysates were incubated with anti-IGF2BP1 or anti-AGO2 antibody and Protein G Dynabeads (Life Technologies) for 30 min at room temperature. After three washing steps with RIP buffer, protein-RNA complexes were eluted by addition of 1% SDS and 65°C for 5 minutes. Protein enrichment was analyzed by western blotting. Co-purified RNAs were extracted using TRIzol and analyzed by RT-qPCR. If indicated, DMSO or 5 µM BTYNB was added 6 h before RIP was performed.

### Western blotting

Infrared western blotting analyses were performed as previously described [20]. Vinculin (VCL/Vcl) served as a loading and normalization control. For RIP analyses VCL served as negative control. For the miTRAP study VCL served as negative control and MS2-BP as control for equal loading of the resin. Antibodies used are indicated in Supplementary Table 6.

### RNA isolation and RT-qPCR

Total RNA from cell lines were isolated using TRIzol. RNA concentration was determined by nanodrop (Tecan Spark). For cDNA synthesis, 2 µg total RNA served as a template using M-MLV Reverse Transcriptase (Promega) and random hexamer primers following manufactureŕs protocol. For RNA decay analyses Oligo-dT primers were used instead of random hexamer primers. RT-qPCR analyses were performed on a LightCycler 480 II (Roche) with the ORA^TM^ qPCR Green ROX L Mix (highQu). In general, RPLP0, EEF2 and GAPDH served as housekeeping genes (normalization controls). For RNA decay analyses RPLP0 served as normalization control. For IGF2BP1 RIP assays RNA data are input normalized and HIST1 served as normalization control. E2F1 served as a positive control. For AGO2 RIP assays RNA data are input normalized and IGF2BP1-KO cells were normalized against control cells. Primer pairs were selected using Primer Blast (https://www.ncbi.nlm.nih.gov/tools/primer-blast/). Sequences are summarized in Supplementary Table 6. Relative RNA abundance was determined by the ΔΔC_t_ method, as previously described [20].

### Nascent RNA capture

For analyzing the newly synthesized RNA the Click-iT^TM^ nascent RNA capture kit (Invitrogen) was used according to manufactureŕs instructions and as previously described [24].

miTRAP experiment. MiTRAP experiments using 3’UTR of MYCN or MS2 control RNA were essentially performed as described recently [46].

### Drug analysis

For determining the EC values 5 x 10^3^ BE(2)-C, 1 x 10^4^ NBL-S or 1.25 x 10^4^ KELLY cells were seeded per well of a 96-well plate. A serial dilution of the drugs was performed and added to the cells at the indicated concentrations. DMSO served as control condition. After 72 h of treatment, an image was acquired using IncuCyte S3 (Sartorius). Confluence was determined using the IncuCyte software. Confluence values were normalized to DMSO control. For the analysis of synergy between two drugs, the cell viability was determined 72 h upon drug exposure using CellTiter GLO (Promega) in a drug matrix screen at indicated concentrations. Synergy relief maps were generated using the SynergyFinder web application (https://synergyfinder.fimm.fi; version 2.0) and the ZIP (Zero interaction potency) method [47]. For BTYNB recovery experiment DMSO or 5 µM BTYNB were added to the cells and grown for 72 h. Afterwards one third of the BTYNB-treated cells were harvested (referred to as ‘3 days BTYNB’) while the rest was treated further with DMSO (‘BTYNB+DMSO 6 days’ = recovery) or 5 µM BTYNB (‘6 days BTYNB’) respectively.

### Animal handling and ethics approval

Animals were handled according to the guidelines of the Martin Luther University. Permission was granted by the state administration office of Saxony-Anhalt (reference number: 42502-2-1381 MLU, 42502-2-1530 MLU, 42502-2-1625 MLU). Female immunodeficient athymic nude mice (FOXN1^nu/nu^) were obtained from Charles River. For subcutaneous xenograft assays 1 x 10^6^ iRFP-labeled BE(2)-C or 2 x 10^6^ iRFP-labeled NBL-S cells (stably transduced using iRFP encoding lentiviruses) were harvested in PBS supplemented with 50% (v/v) matrigel (Sigma) and injected into the left flank of six-week old mice. Prior injection, cells were counted using a TC20 Cell Counter (Bio-Rad). Subcutaneous tumor growth and volume were measured and monitored by non-invasive near-infrared imaging using a Pearl Impulse Imaging System (LI-COR) and isoflurane as anesthetic. Tumor volume was calculated using the formula π/6 × L1 × L2 × L3. For testing BTYNB *in vivo* BE(2)-C cells were treated for 24 h with BTYNB or DMSO prior to harvesting. For intra-tumoral application of BTYNB 5 x 10^5^ iRFP-labeled BE(2)-C cells were injected into nude mice and grown for 2 weeks. Afterwards treatment was started by injecting 50 mg/kg body weight BTYNB in 100 µl 35% cyclodextrin solution in PBS with a DMSO concentration of 7%. Treatment was performed five days in a row followed by two days without compound application before the cycle started again. Transgenic mice were generated by Taconic Bioscience except of LSL- *MYCN* mice which were obtained from Prof. Schulte, Berlin. For getting a homogenous background, mice were backcrossed to C57Bl6/N mice (Charles River) and afterwards crossbred. Mouse genotyping was confirmed by PCR. Mice were palpated weekly for abdominal tumors. Mice are summarized in Supplementary Table 7. Primer sequences are provided in Supplementary Table 6. Patient-derived xenografts (PDX) were performed by EPO company in Berlin, Germany. The neuroblastoma tumor 14647 harboring *MYCN* amplification and 17q gain was implemented in NOG mice to generate PDX. Treatment was performed by intra-peritoneal application of DMSO, 100 mg/kg body weight BTYNB, 2.5 mg/kg body weight YM-155 or the combination of BTYNB and YM-155 in 100 µl 30% cyclodextrin solution in PBS with a final DMSO concentration of 7%. Compound application was done five days in a row followed by two days without treatment. Tumor volume and mouse weight was documented over time.

### BTYNB synthesis and stability

BTYNB was synthesized starting from 5-Bromo-2- thiophenecarboxaldehyde and anthranilamide. In detail, a mixture of 5-Bromothiophene-2- carboxaldehyde (1 mmol) and anthranilamide (1 mmol) in ethanol (5 ml) was refluxed for 3 hours then cooled to room temperature. The precipitated solid was then filtered, washed with ethanol and hexane and air dried. Yield: 70%; Analytical characterization: MS m/z: 310.10 [M+H]+; 1H NMR (400 MHz, DMSO) δ 8.49 (s, 1H), 7.60 (dd, J = 7.7, 1.3 Hz, 1H), 7.31 – 7.22 (m, 2H), 7.06 (d, J = 3.8 Hz, 1H), 6.93 (d, J = 3.8 Hz, 1H), 6.78 – 6.64 (m, 2H), 5.95 (t, J = 2.2 Hz, 1H); HPLC: rt 11.31 min (Purity of BTYNB 97.77 %) Chemical stability of BTYNB. A 10 µM BTYNB solution in acetonitrile was incubated with Dulbecco’s Modified Eagle Medium (DMEM) at 37 °C, and samples were analyzed at different time intervals (0 min, 30 min, 60 min, 120 min, 24 h, 48 h and 72 h) using the HPLC method described below to determine and quantify potential degradation products. A good stability profile up to 72 h was obtained (Table 1; Supplementary Fig. 6e). In addition this protocol was used to test the stability of BTYNB under acidic conditions where the solution was incubated with 10% v/v of triflouroacetic acid in DMSO at 37 °C. A slight degradation of 15 % after 72 h was observed (Table 1; Supplementary Fig. 6f).

**Table 1:** BTYNB stability

| condition | 0h | 6h | 12h | 24h | 48h | 72h |
| --- | --- | --- | --- | --- | --- | --- |
| **BTYNB in DMEM** | 100% | 100.4 % | 101.2 % | 103.1 % | 105.3 % | 107.7 % |
| **BTYNB in DMEM under acidic condition** | 100% | 98.3 % | 97.4 % | 95.1 % | 85.8 % | 84.3 % |

### HPLC method

As HPLC system a LiChrosorb® RP-18 (5 µm) 100-4.6 column from the manufacturer Merck, two LC-10AD pumps, a SPD-M10A VP PDA detector, and a SIL-HT autosampler were used (all from the manufacturer Shimadzu, Kyoto, Japan). As mobile phase a gradient with increasing polarity composed of methanol/water/triflouroacetic acid at a flow rate of 1 ml/minute was used. UV absorbance was measured at 254 nm.

Plasma protein binding of BTYNB. Serial dilutions of BTYNB in acetonitrile (0-100 µM) were prepared and their absorbance was measured by the described HPLC method to get a calibration curve. Next, a 100 µM solution of BTYNB in acetonitrile was incubated with fetal bovine serum (FBS) or human serum albumin (HSA; 40 mg/ml in PBS) and samples at different time intervals (0 min, 30 min, 60 min, 120 min, 24 h, 48 h and 72 h) were analyzed. The samples were subjected to ultra- centrifugation (5 min, 10000 rpm/g) using modified PES 30K low protein binding centrifugal filter. The filtrate was then analyzed using HPLC to determine the available concentration of BTYNB (using the previous calibration curve) and also to determine and quantify other potential degradation products. The obtained results showed that BTYNB has a very high plasma protein binding upon testing either in FBS or HAS (Supplementary Fig. 6g, h).

### Human neuroblastoma samples

Human neuroblastoma cohort consists of 69 previously published untreated primary samples [48] extended by 31 additional tumor specimens. Tumors were granted after patient consent and ethical approval from the Cologne tumor bank and the Universitätsklinikum Essen, Germany (Supplementary Table 8). The International Neuroblastoma Staging System criteria (INSS) was used for staging.

### Tissue preparation

Transgenic tumors and murine organs were dissected and shock-frozen on dry ice. Approximately 30 mg of frozen tissue was lysed, homogenized with Zirconium Oxide beads and Precellys 24 homogenizer (berting technologies) and lysates were subjected to the Qiagen AllPrep DNA/RNA/Protein Kit protocol, with the exception, that precipitated protein was solubilized in 5% SDS solution. Human tumor samples were handled as previously described [48].

### Immunohistochemistry

Transgenic tumor samples were fixated with Roti®-Histofix (Roth) and embedded in paraffin. Paraffin blocks were sliced at 4 µm thickness using LEICA RM2235 (Leica) and transferred to a specimen slide. Staning was performed for hematoxylin and eosin as well as Phox2b.

Images were acquired using a Nikon Eclipse TE2000-E microscope with a 20x objective and 1.5 manual magnification.

### Plasmids and cloning

Cloning strategies including vectors, oligonucleotides used for PCR and restrictions sites are summarized in Supplementary Table 6. All constructs were validated by sequencing.

### Statistics and presentation

All experiments were at least performed in biological triplicates. Western blotting and genotyping PCR for transgenic mouse experiments were just performed once as in this case biological replicates are individual mice. The western blot for the miTRAP experiment was also done once as we performed four experiments in total and three off them went completely for miRNA sequencing. Statistical significance was tested by parametric two-sided Student’s *t*-test on equally distributed data (errors defined as standard deviation of the mean). This includes all lab-based analyses (western blot, RT-qPCR, drug assays, luciferase assays, viability measurements). Otherwise, a non-parametric two-sided Mann-Whitney-test was performed (error defined as standard error of the mean). This includes all tumor-related analyses (human neuroblastoma samples, transgenic mouse and xenograft data). For box plots, horizontal lines demonstrate the median with upper and lower box boundaries demonstrating the 25^th^-75^th^ centiles. Error bars represent the maximum and minimum (Fig. 3g, h; Supplementary Fig. 6d). For Kaplan-Meier survival analyses, statistical significance was determined by log-ranked test. No testing for outliers were performed. *, p < 0.05; **, p < 0.01; ***, p < 0.001; ****, p < 0.0001. Data were visualized with GraphPad Prism (version 8.0.1). Heatmaps were generated with Flourish web application (https://flourish.studio/). Biorender (https://www.biorender.com/) was used for generation of some figures.

## Results

### MNA neuroblastoma essential genes are enriched on chromosome 17q

Gene expression and genomic aberrations in neuroblastoma were analyzed in 100 tumors, representing all INSS stages (Supplementary Table 8). Shallow whole genome sequencing (sWGS) identified common chromosomal gains and deletions (Fig. 1a; Table 2) [1, 2]. The most frequent disturbances were gains at Chr 17 with 35 tumors showing uneven copy number of 17q and 17p (referred to as unbalanced 17q).

**Table 2:** Chromosomal aberrations in neuroblastoma

| chromosomal region | genomic aberration | frequency | frequency literature | references |
| --- | --- | --- | --- | --- |
| 1p | loss | 10% | 30-40% | [2, 49] |
| 2p | gain | 23% | 20-35% | [2, 49] |
| 3p | loss | 24% | 18% | [50] |
| 4p | loss | 14% | 20% | [2] |
| 7 | gain | 44% | 40% | [51] |
| 11q | loss | 27% | 29-44% | [2, 49, 50] |
| 14q | loss | 20% | 22-25% | [2, 49] |
| 17q | gain | 79% | > 50% | [2, 49] |
| unbalanced 17q | gain | 35% | - |  |

Survival analyses demonstrated association of unbalanced 17q with poor survival (Fig. 1b). Adverse prognosis was fostered by concomitant *MYCN* amplification (Fig. 1b). Reduced survival probability was confirmed for the 17 MNA tumors compared to *MYCN* non-amplified (nMNA) tumors (Fig. 1c). Only few tumors (n = 4) with balanced 17q and MNA were observed prohibiting conclusive survival analyses (Supplementary Fig. 1a). This suggested pro-oncogenic association of 2p gain and unbalanced 17q in neuroblastoma.

To identify essential candidate genes, we analyzed gene dependency scores via the DepMap portal [38] for 13 MNA neuroblastoma and 607 non-neuroblastoma cell lines. This revealed 177 genes (175 autosomal, 2 gonosomal) with increased essentiality in MNA neuroblastoma. *MYCN*, *ISL1* and *HAND2* were the top-ranked genes (Supplementary Table 2). Only for Chr 17, significant enrichment of essential genes, with top ranking of *IGF2BP1*, *GJC1* and *MSI2,* was observed (Fig. 1d, e; Table 3).

**Table 3:** Hypergeometric testing for distribution of neuroblastoma essential genes

| Chr | protein coding genes | NB essential genes | % distribution | p-value |
| --- | --- | --- | --- | --- |
| 1 | 2048 | 22 | 12.6% | 0.2496822 |
| 2 | 1247 | 17 | 9.7% | 0.0668034 |
| 3 | 1075 | 4 | 2.3% | 0.9904822 |
| 4 | 751 | 5 | 2.9% | 0.8245106 |
| 5 | 886 | 4 | 2.3% | 0.9651801 |
| 6 | 1047 | 6 | 3.4% | 0.9238733 |
| 7 | 917 | 4 | 2.3% | 0.9716455 |
| 8 | 683 | 3 | 1.7% | 0.9527968 |
| 9 | 781 | 5 | 2.9% | 0.8492704 |
| 10 | 731 | 4 | 2.3% | 0.9076264 |
| 11 | 1311 | 6 | 3.4% | 0.9835301 |
| 12 | 1035 | 10 | 5.7% | 0.4815816 |
| 13 | 321 | 3 | 1.7% | 0.5682151 |
| 14 | 612 | 0 | 0.0% | 1 |
| 15 | 599 | 7 | 4.0% | 0.3128974 |
| 16 | 853 | 5 | 2.9% | 0.8971414 |
| **17** | **1188** | **52** | **29.7%** | **5.047e-22** |
| 18 | 268 | 3 | 1.7% | 0.4479059 |
| 19 | 1471 | 9 | 5.1% | 0.9305177 |
| 20 | 546 | 2 | 1.1% | 0.9626367 |
| 21 | 232 | 0 | 0.0% | 1 |
| 22 | 444 | 4 | 2.3% | 0.5853077 |

Chr – chromosome; NB – neuroblastoma

This further supports the previously proposed prognostic value of IGF2BP1 [14] in neuroblastoma, as observed here in low- and high-risk subgroups except MNA (Fig. 1f; Supplementary Fig. 1b-f). In agreement, unbalanced 17q is associated with adverse outcome in nMNA tumors (Supplementary Fig. 1g, h). Cox multi-variate analyses validated MNA, unbalanced 17q as well as IGF2BP1 and MYCN expression as independent predictors of poor outcome (Table 4).

**Table 4:**
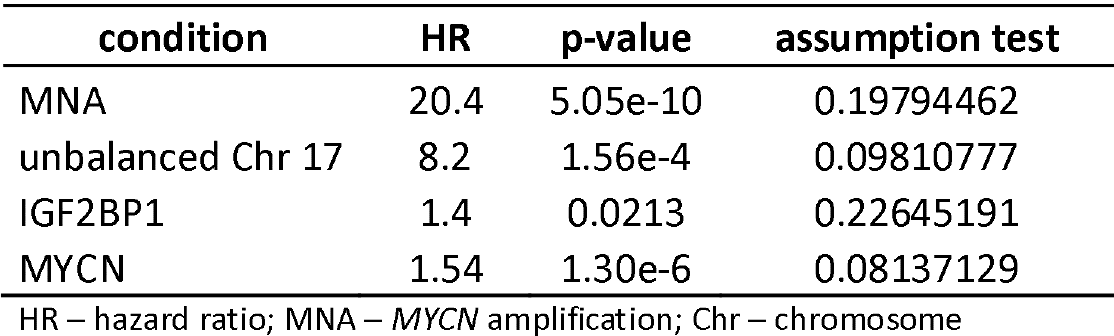
Cox multi-variate analyses

| condition | HR | p-value | assumption test |
| --- | --- | --- | --- |
| MNA | 20.4 | 5.05e-10 | 0.19794462 |
| unbalanced Chr 17 | 8.2 | 1.56e-4 | 0.09810777 |
| IGF2BP1 | 1.4 | 0.0213 | 0.22645191 |
| MYCN | 1.54 | 1.30e-6 | 0.08137129 |

HR – hazard ratio; MNA – MYCN amplification; Chr – chromosome

However, IGF2BP1 showed strong association with MYCN and was significantly increased in MNA, unbalanced 17q, INSS-4 and deceased patients (Supplementary Fig. 1i-m). Correlation analyses in two independent tumor cohorts showed substantial relation of IGF2BP1- and MYCN-associated gene expression (Supplementary Fig. 1n). This was less pronounced for other transcriptional regulators like the neuroblastoma core regulatory circuit [52] member PHOX2B (Table 5).

**Table 5:** Correlation of IGF2BP1- and MYCN-associated gene expression

| gene | Bell |  | Kocak |  |
| --- | --- | --- | --- | --- |
|  | R Spearman | p value | R Spearman | p value |
| **MYCN** | 0,9286 | < 0,0001 | 0,7306 | < 0,0001 |
| **PHOX2B** | 0,7359 | < 0,0001 | 0,4511 | < 0,0001 |
| **GATA3** | 0,7351 | < 0,0001 | 0,5584 | < 0,0001 |
| **HAND2** | 0,7193 | < 0,0001 | 0,2191 | < 0,0001 |
| **SOX4** | 0,6696 | < 0,0001 | 0,0270 | 0,0005 |
| **NFkB** | 0,5332 | < 0,0001 | - 0,4308 | < 0,0001 |
| **PRRX1** | - 0,1866 | < 0,0001 | - 0,1979 | < 0,0001 |

Depletion studies in MNA and 17q unbalanced BE(2)-C cells confirmed association of deregulated gene expression upon MYCN and IGF2BP1 knockdown (Supplementary Fig. 1o). Public neuroblastoma single cell RNA-seq data [35] indicated nearly exclusive expression and strong association of IGF2BP1 and MYCN expression in tumor cell populations (Supplementary Fig. 1p, q). Collectively, this suggested an interconnected disease driving role of IGF2BP1 and MYCN in HRN.

### IGF2BP1 is a post-transcriptional enhancer of MYCN expression

We demonstrated that IGF2BP1 modulates MYCN expression in neuroblastoma cell models [14, 53], but the underlying mechanisms and potential synergy of both in promoting HRN remained elusive. IGF2BP1-CLIP studies in hESCs indicated association of IGF2BP1 at the MYCN 3’UTR (Supplementary Fig. 2a). The deletion (KO) of IGF2BP1 consistently reduced MYCN protein and mRNA expression in three neuroblastoma cell lines (Fig. 2a; Supplementary Fig. 2b). In line with IGF2BP1’s main role in cancer [20, 21, 54], IGF2BP1-KO significantly decreased MYCN mRNA half-life in BE(2)-C (Fig. 2b). IGF2BP1-RIP confirmed pronounced association of IGF2BP1 with the validated target mRNA E2F1 [24] as well as MYCN transcripts (Fig. 2c; Supplementary Fig. 2c, 3a). Recent miTRAP (microRNA trapping by RNA affinity purification [46]) studies indicated miRNA and AGO2 association with the MYCN 3’UTR in BE(2)-C [53]. Likewise, miTRAP demonstrated association of IGF2BP1 with the MYCN 3’UTR (Fig. 2d). This suggested that MYCN expression is druggable by BTYNB, a small molecule inhibitor of

**Fig. 2:**
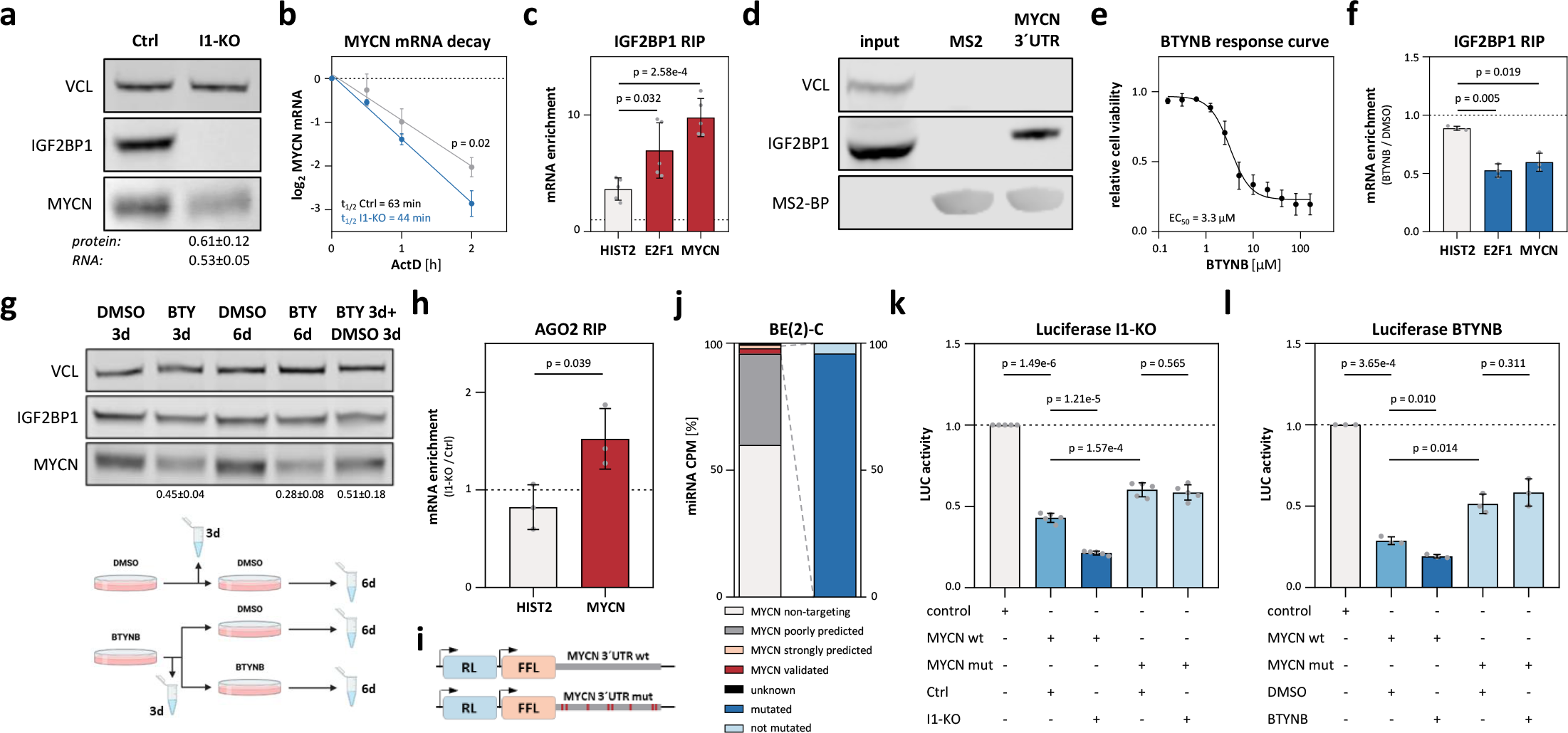
IGF2BP1 stimulated MYCN expression is 3’UTR-, miRNA- and BTYNB-dependent. **(a)** Western blot (n = 6) and RT-qPCR (n = 5) analysis of MYCN expression upon IGF2BP1-KO (I1-KO) in BE(2)-C. **(b)** MYCN mRNA decay monitored by RT-qPCR in control (grey) and I1-KO (blue) BE(2)-C upon indicated time of actinomycin D (ActD) treatment (n = 3). **(c)** IGF2BP1-RIP analyses in parental BE(2)-C (n = 5). **(d)** Western blotting of miTRAP from BE(2)-C lysates using MS2-fused MYCN 3’UTR or MS2-RNA (n = 1). **(e)** BTYNB-response curve in BE(2)-C (n = 3). **(f)** IGF2BP1-RIP in BE(2)-C treated with DMSO or BTYNB (n = 3). **(g)** Western blot analysis of MYCN expression in BE(2)-C treated with BTYNB (BTY, n = 3) for 3 and 6 d or replacement of BTY by DMSO after 3 d treatment. Treatment scheme in lower panel. **(h)** AGO2-RIP analyses in I1-KO versus control BE(2)-C (n = 3). **(i)** Scheme of luciferase reporters constructs. **(j)** Expression of miRNAs in BE(2)-C (left) and mutated fraction of MYCN-targeting miRNA-binding sites (right). (k, l) Activity of indicated luciferase reporters in control and I1- KO (k, n = 5) or DMSO- and BTYNB-treated BE(2)-C (l, n = 3).

IGF2BP1-mRNA association impairing IGF2BP1-driven E2F1 and MYC expression [24]. BTYNB decreased neuroblastoma cell viability, as previously reported [55], disturbed MYCN mRNA association with IGF2BP1 and impaired MYCN at only modest reduction of IGF2BP1 (Fig. 2e-g; Supplementary Fig. 2d, e, 3b). MYCN decrease was pronounced by extended treatment and recovered after compound withdrawal, indicating that BTYNB reversibly impairs IGF2BP1-stimulated expression of MYCN.

**Fig. 3:**
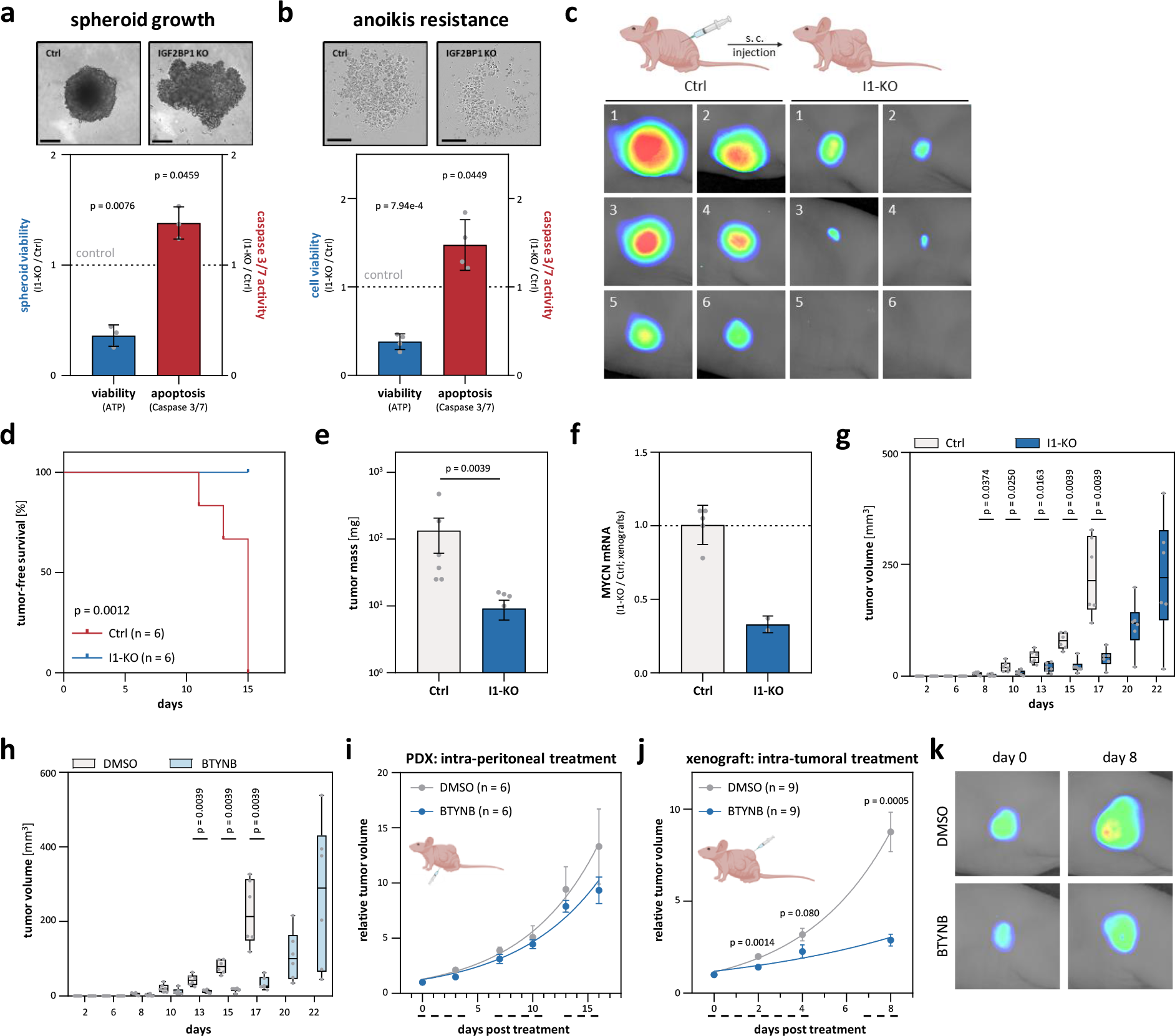
IGF2BP1 deletion and inhibition by BTNYB impairs xenograft tumor growth. (a, b) The viability and caspase3/7-activity of parental (Ctrl) and I1-KO BE(2)-C was analyzed in spheroid growth (a, n = 3) and anoikis-resistance studies (b, n = 4; bars a, 200 µm and b, 400 µm). (c-e) Tumor growth (n = 6) of Ctrl and I1-KO BE(2)-C s.c. xenografts was monitored by non-invasive infrared imaging **(c)**, tumor-free survival **(d)** and final tumor mass **(e)**. **(f)** RT-qPCR analysis of MYCN mRNA levels in excised xenograft tumors and non-palpable tumor cell mass upon I1-KO (Ctrl, n = 5; I1-KO, n = 2). (g, h) Tumor growth (n = 6) of Ctrl and I1-KO **(g)** or DMSO- and BTYNB-pretreated **(h)** BE(2)-C xenografts was monitored by tumor volume over time. **(i)** Tumor growth (n = 6) was monitored by relative volume of s.c. PDX tumors treated i.p. with DMSO (grey) or BTYNB (blue). Daily treatment in three cycles is indicated by dashed lines below the x-axis. **(j)** Tumor growth (n = 9) was monitored by relative volume of s.c. BE(2)-C treated i.t. with DMSO (grey) or 50 mg/kg BW BTYNB (blue). Daily treatment in two cycles is indicated by dashed lines below the x-axis. **(k)** Representative images of tumors at start (day 0) and final treatment (day 8).

Whether IGF2BP1-dependent regulation of MYCN is miRNA-dependent was investigated by AGO2-RIP. IGF2BP1-KO significantly increased association of AGO2 with the MYCN mRNA (Fig. 2h; Supplementary Fig. 3c). This was analyzed further by luciferase reporters lacking a native 3’UTR (control) and comprising the wildtype (wt) or mutated (mut) MYCN 3’UTR. In the latter, >90% of predicted/validated miRNA targeting sites were inactivated by mutation (Fig. 2i, j; Table 6).

**Table 6:** Validated and predicted miRNA-binding sites for MYCN

| miRNA family | miRNA binding | binding site<br>(MYCN 3'UTR bp) | luciferase reporter |
| --- | --- | --- | --- |
| miR-34-5p | validated | 22-29 | mutated |
| miR-19-3p | validated | 31-38 | mutated |
| miR-29-3p | validated | 332-339 | mutated |
| miR-101-3p | validated | 493-500 | mutated |
| let-7-5p/miR-98-5p | validated | 505-511 | mutated |
| miR-101-3p | validated | 562-568 | wildtype |
| miR-34-5p | validated | 579-585 | mutated |
| miR-302-3p | predicted | 858-864 | mutated |
| miR-17-5p/20-5p/93-5p/106-5p | predicted | 859-865 | mutated |
| let-7-5p/miR-98-5p | validated | 868-874 | mutated |

miRNA – microRNA; 3'UTR – 3' untranslated region; bp – base pairs

Activity of the wt reporter was consistently reduced in comparison to control and mut reporters, confirming conserved miRNA-dependent regulation of the MYCN 3’UTR in neuroblastoma cell models (Fig. 2k; Supplementary Fig. 2f, g). Both, IGF2BP1-KO and BTYNB, further decreased wt reporter activity, whereas the mut reporter remained unaffected (Fig. 2l; Supplementary Fig. 2h). This indicated that BTYNB disrupts miRNA-dependent regulation of MYCN by IGF2BP1.

IGF2BP1-dependent stabilization of some target mRNAs, e.g. MYC and E2F1, is enhanced by m^6^A modification via the METTL3/14 methyltransferase complex [24]. To test if this is also observed for the m^6^A-modified MYCN mRNA (Supplementary Fig. 4a), was analyzed by METTL3/14 co-depletion. Knockdown reduced E2F1 protein and mRNA abundance without affecting IGF2BP1 expression (Supplementary Fig. 4b-d), as prior reported [24]. However, MYCN protein and mRNA levels remained unchanged, suggesting that IGF2BP1 promotes MYCN expression largely independent of m^6^A, but in a 3’UTR-, miRNA- and BTYNB-dependent manner.

### IGF2BP1 is an essential and druggable driver of neuroblastoma growth

Our findings suggested, IGF2BP1 as a novel driver of neuroblastoma. IGF2BP1-KO consistently decreased spheroid growth and anoikis resistance, whereas caspase 3/7 activity (apoptosis) was moderately increased (Fig. 3a, b; Supplementary Fig. 5a, b). In subcutaneous (s.c.) xenografts, IGF2BP1-KO impaired tumor engraftment, MYCN expression and delayed growth by 5-7 days (Fig. 3c- g; Supplementary Fig. 5c-f). This was also observed by BTYNB pre-treatment of parental cells (Fig. 3h). How the intra-peritoneal (i.p.) delivery of BTYNB affects growth of an MNA and 17q unbalanced s.c. patient-derived xenograft (PDX) was analyzed by three cycles of treatment at 100 mg/kg body weight (BW). Notably, treatment showed no obvious toxicity (Supplementary Fig. 6a). Tumor growth was only slightly reduced by BTYNB (Fig. 3i). This rather moderate potency is probably due to high plasma protein binding of BTYNB, suggesting poor pharmacokinetics (Supplementary Fig. 6e-h). Aiming to overcome these limitations, s.c. BE(2)-C xenograft tumors were treated with two cycles of intra-tumoral (i.t.) BTYNB application at 50 mg/kg BW (Fig. 3j; Supplementary Fig. 6b-d). As for i.p. treatment, no obvious signs of toxicity were observed. However, i.t. treatment substantially impaired tumor growth, indicating strong target potential of IGF2BP1 and therapeutic lead prospects of BTYNB.

### MYCN and IGF2BP1 form a druggable positive feedforward loop in neuroblastoma

MYC promotes IGF2BP1 transcription [56]. In view of MYC/N similarity [57] and strongly associated expression of IGF2BP1 and MYCN in neuroblastoma (Supplementary Fig. 1i), we hypothesized that MYCN promotes IGF2BP1 transcription as well. Public MYCN ChIP-seq data [44] indicated conserved association of MYCN at the *IGF2BP1* promoter and coverage of several E-Box motifs (Fig. 4a). MYCN depletion substantially reduced steady state level and nascent transcript synthesis of IGF2BP1, indicating MYCN-driven IGF2BP1 transcription and feedforward regulation of MYCN/IGF2BP1 (Fig. 4b, c; Supplementary Fig. 7a). To test if this is also druggable via MYCN, we explored four bromodomain inhibitors (BRDi: Mivebresib, ARV-771, CPI-0610, INCB-057643), which impair MYC/N abundance and transcriptional activity (Fig. 4d) [58, 59]. In MNA cells, BRDi concisely decreased IGF2BP1 and MYCN expression with the most robust downregulation observed by Mivebresib (Fig. 4e; Supplementary Fig. 7b, c). In contrast, MYCN and IGF2BP1 expression remained unaffected by BRDi in nMNA NBL-S, in which MYCN synthesis is elevated by genomic translocation (Supplementary Fig. 7c) [60]. In view of MYCN/IGF2BP1 feedforward regulation, it was expected that IGF2BP1 alters sensitivity to BRDi and BTYNB (IGF2BP1i) synergizes with BRDi in non-MYCN-translocated neuroblastoma. IGF2BP1-KO severely reduced EC_50_ values for Mivebresib (∼7-fold) and drug matrix screens confirmed substantial benefit of combined treatment (maximum synergy score = 19-21) exclusively in MNA cells (Fig. 4f, g; Supplementary Fig. 7d-f). This suggested effective targeting of MYCN/IGF2BP1 by combined BRDi and IGF2BP1i in MNA neuroblastoma.

**Fig. 4:**
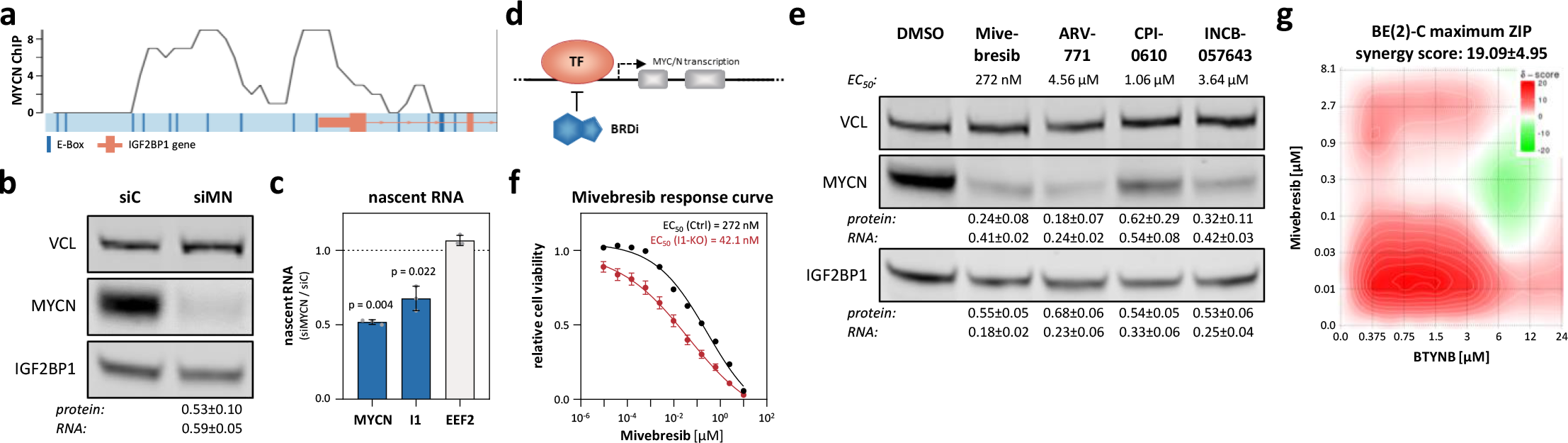
MYCN-driven IGF2BP1 expression is impaired by BRD inhibitors. **(a)** MYCN ChIP-seq profile of the IGF2BP1 promoter region. E-Boxes, putative MYC/N-binding sites, are indicated in dark blue. The IGF2BP1 gene is depicted schematically in orange up to the beginning of the second intron. **(b)** Western blot (n = 3) and RT-qPCR (n = 6) analysis of IGF2BP1 expression upon MYCN (siMN) compared to control knockdown (siC) in BE(2)-C. **(c)** RT-qPCR (n = 3) analysis of indicated nascent mRNAs upon MYCN compared to control knockdown in BE(2)-C (I1 - IGF2BP1). **(d)** Scheme of putative MYCN regulation by BRD (TF - transcription factor). **(e)** Western blot and RT-qPCR analysis of MYCN and IGF2BP1 expression upon treatment of BE(2)-C with indicated BRD inhibitors (n = 3). **(f)** Mivebresib response curve in control (black) and I1-KO (red) BE(2)-C (n = 4). **(g)** Relief plot depicting the ZIP synergy for combined treatment of BTYNB and Mivebresib in BE(2)-C (n = 3).

### IGF2BP1/MYC(N)-driven genes on Chr 17q harbor therapeutic potential in cancer treatment

Feedforward regulation by MYCN/IGF2BP1 likely impact many effectors due to broad target spectra [57, 61], suggesting MYCN/IGF2BP1 also synergize with oncogenic genes located on Chr 17q. This was evaluated for selected oncogenes/tumor suppressors (control) and essential genes located at Chr 17q by integrating: i) gene dependency score in MNA neuroblastoma cell lines, ii) fold change of mRNA in 17q unbalanced, MNA or INSS-4 tumors, iii) gene hazardous ratio, iv) correlation of mRNA expression with MYCN and IGF2BP1 in tumors, v) fold change of mRNA upon MYCN and IGF2BP1 depletion, vi) MYCN ChIP-seq data, vii) IGF2BP1-CLIP data (Supplementary Fig. 8a; Supplementary Tables 2, 4, 5, 9). Next to *IGF2BP1*, this analysis unraveled top-ranking of *BIRC5* and *TOP2A* among common essential as well as *NME1* among neuroblastoma essential Chr 17q genes. This was further evaluated for the oncogene BIRC5, which was proposed for targeted treatment (BIRC5i) of various cancers including HRN (Supplementary Fig. 8b-g) [26, 62–64]. Regulation by MYCN/IGF2BP1 was confirmed by: i) BIRC5 downregulation upon MYCN and IGF2BP1 depletion, ii) ChIP-seq data indicating conserved MYCN association at the *BIRC5* promoter, iii) IGF2BP1-CLIP demonstrating conserved IGF2BP1 association at the BIRC5 3’UTR, iv) reduced activity of BIRC5 3’UTR luciferase reporters upon IGF2BP1-KO and IGF2BP1i by BTYNB (Fig. 5a-c; Supplementary Fig. 8h, i). Like observed for BRDi, IGF2BP1-KO reduced EC_50_ values (∼3-fold) for BIRC5i by YM-155 (Fig. 5d), which was well tolerated in clinical trials and showed anti-tumor activity in combination therapy [26, 63, 64]. Accordingly, we investigated the response of MNA/17q unbalanced s.c. PDX tumors to YM-155 (2.5 mg/kg BW) and combined i.p. treatment with BTYNB (100 mg/kg BW). Although results remained non-significant, YM-155 modestly reduced tumor growth and this was further enhanced in combination with BTYNB (Fig. 5e). Drug matrix analyses in MNA cells supported a moderate benefit of combined BTYNB/YM-155 treatment (Fig. 5f; Supplementary Fig. 8j). In sum, this suggested that IGF2BP1 is a potent, druggable oncogene in neuroblastoma synergizing with MYCN in a transcriptional/post-transcriptional feed-forward loop resulting in the upregulation of oncogenes like the 17q-located BIRC5 (Fig. 5g).

**Fig. 5:**
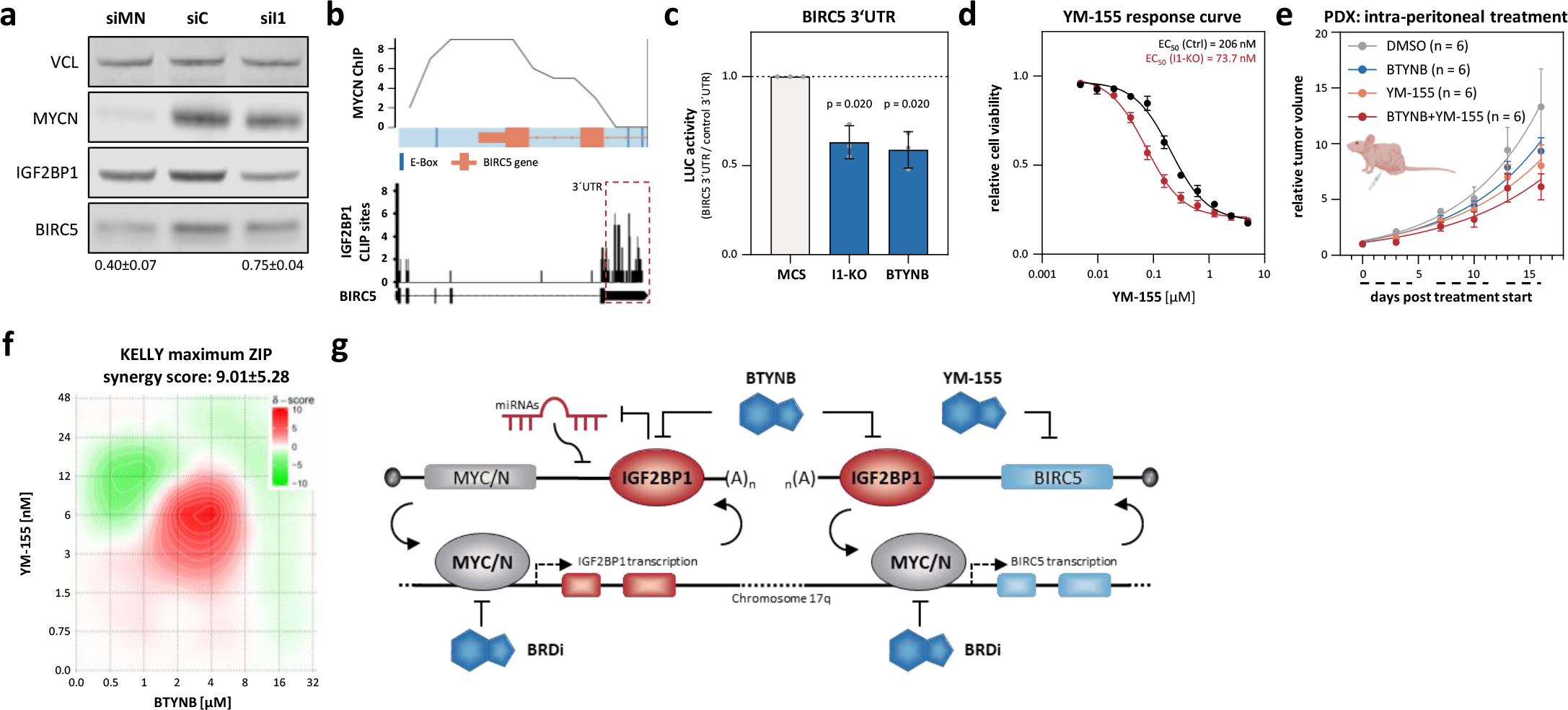
MYCN/IGF2BP1 feedforward regulation promotes expression of druggable oncogenes located on Chr 17q. **(a)** Western blot analysis of BIRC5 expression upon MYCN (siMN) or IGF2BP1 (siI1) compared to control (siC) knockdown in BE(2)-C (n = 3). **(b)** MYCN ChIP-seq profile of the BIRC5 promoter region (upper panel) and IGF2BP1-CLIP profile for the BIRC5 mRNA (lower panel). **(c)** BIRC5 3’UTR luciferase activity in I1-KO versus parental (Ctrl) and BTYNB- versus DMSO-treated BE(2)-C (n = 3). **(d)** YM-155 response curve in control (black) and I1-KO (red) BE(2)-C (n = 4). **(e)** Tumor growth (n = 6) was monitored by relative volume of s.c. PDX tumors treated i.p. with DMSO (grey), YM-155 (orange) or in combination of YM-155 and BTYNB (red). Daily treatment in three cycles is indicated by dashed lines below the x-axis. **(f)** Relief plot showing the ZIP synergy for combined treatment of BTYNB and YM-155 in KELLY (n = 3). **(g)** Scheme of transcriptional/post-transcriptional feedback regulation of MYC/N, IGF2BP1 and downstream effectors as well as treatment possibilities.

### IGF2BP1 is a strong oncogenic driver of neuroblastoma and promotes MYCN protein stability

Aiming to test IGF2BP1’s oncogenic potential in a sympatho-adrenal neuroblastoma model, we established transgenic LSL-*IGF2BP1*-IRES-iRFP mice (Supplementary Fig. 9a), essentially as previously described for MYCN and Lin28b [65, 66]. Notably, the ∼9 kb IGF2BP1 3’UTR was discarded to enhance expression by preventing miRNA-directed downregulation, primarily by let-7 miRNAs [22]. Conditional transgene expression in adrenal glands and peripheral nerves was induced by *Dbh*-driven iCRE. Heterozygous or homozygous transgene integration at the *Rosa26* (*R26*) locus was validated for all analyzed animals by gDNA-PCR (Supplementary Fig. 9-12). LSL-*IGF2BP1* mice were fertile without any obvious phenotypes and offspring were born according to Mendelian ratio. No tumors were observed in controls, LSL-*IGF2BP1* or *Dbh*-iCRE (data not shown). Heterozygous R26^IGF2BP1/-^ mice only showed low tumor burden (1/8) within one year (Fig. 6a). In sharp contrast, homozygous R26^IGF2BP1/IGF2BP1^ mice developed tumors with an incidence of 100% (7/7; Fig. 6a). Ethical culling was required after 169-305 d (median survival 234 d), suggesting a strongly IGF2BP1-dose dependent tumor induction. Seven out of eight IGF2BP1-induced tumors resided in the lower abdomen, potentially arising from sympathetic nerves, and only one homozygous tumor was derived from the adrenal gland (Fig. 6b). In contrast to other neuroblastoma models, no other primary tumor sites, like the superior cervical ganglia, were observed [65–67]. Surprisingly, no tumors (0/6) were found in the *Dbh*-driven heterozygous R26^MYCN/-^ model. Strikingly, however, tumor incidence increased to 100% (8/8) and median survival was reduced to 107 d in R26^IGF2BP1/MYCN^ mice, validating a strong synergy of IGF2BP1 and MYCN in promoting neuroblastoma (Fig. 6a). Moreover, R26^IGF2BP1/MYCN^ substantially elevated tumor burden per animal with 19 tumors observed in 8 animals. Of these, 75% harbored tumors derived from adrenal glands, 62.5% contained abdominal tumors and 50% showed tumors along the spine (Fig. 6b). Inspection of tumors indicated neuroblastoma characteristic small round blue tumor cell morphology and elevated nuclear Phox2b protein (Fig. 6c). Principal component analyses of RNA-seq data derived from tumors and/or adrenal glands revealed no striking differences in respect to tumor location or between R26^IGF2BP1^ and R26^IGF2BP1/MYCN^ tumors (Fig. 6d; Supplementary Fig. 9c). The investigation of adrenergic (ADRN) versus mesenchymal-like (MES) gene expression signatures [33], suggested mesenchymal-like pattern in wildtype (AG) and R26^MYCN/-^ (AGM) adrenal glands (Fig. 6e). In sharp contrast, rather adrenergic neuroblastoma was observed in both, R26^IGF2BP1^ (TI) and R26^IGF2BP1/MYCN^ (TIM) tumors. In support of MYCN/IGF2BP1-driven expression of 17q oncogenes, e.g. BIRC5, TI and TIM samples showed elevated expression of Mycn, Igf2bp1 and Birc5, whereas Myc was diminished (Fig. 6f). This was associated with elevated expression of neuroblastoma (e.g. Phox2b), sympatho-adrenal (e.g. Th), neural (e.g. Gap43), neural stem cell (e.g. Msi1) and neural crest cell markers (e.g. Tfap2b) [65, 68]. At the protein level, transgene dose-dependent upregulation of IGF2BP1/Igf2bp1 was observed in non-tumorous adrenal glands and further fostered in tumors (Fig. 6g). The investigation of MYCN/Mycn protein abundance in adrenal glands and tumors revealed a striking IGF2BP1-dependent increase without obvious association of altered mRNA levels (Fig. 6h). This suggested IGF2BP1-dependent modulation of MYCN protein turnover (Fig. 6i), since the *MYCN* transgene lacks the native 3’UTR. In support of this, IGF2BP1 depletion decreased MYCN protein levels in TET21N, lacking the native MYCN 3’UTR (Fig. 6j) [69]. In emetine treated TET21N, MYCN protein showed an approximately 2-fold decreased half-life upon IGF2BP1 knockdown (Fig. 6k; Supplementary Fig. 13a), indicating that IGF2BP1 promotes both, MYCN mRNA and protein stability. Investigation of key regulators controlling MYCN protein turnover [70], revealed upregulation of stabilizing factors (Aurka, Plk1, Cdk1) especially in TI (Fig. 6l). Their abundance was decreased by IGF2BP1 depletion in human neuroblastoma cells without striking effects on post-translational modifications (Fig. 6m, n; Supplementary Fig. 13b). In contrast, no obvious association was observed for IGF2BP1 and factors destabilizing MYCN protein, primarily GSK3B/Gsk3b and FBXW7/Fbxw7. This indicated that IGF2BP1 promotes MYCN protein stability, probably by the previously described RNA-dependent enhancement of AURKA [54] and additional upregulation of PLK1 and CDK1.

**Fig. 6:**
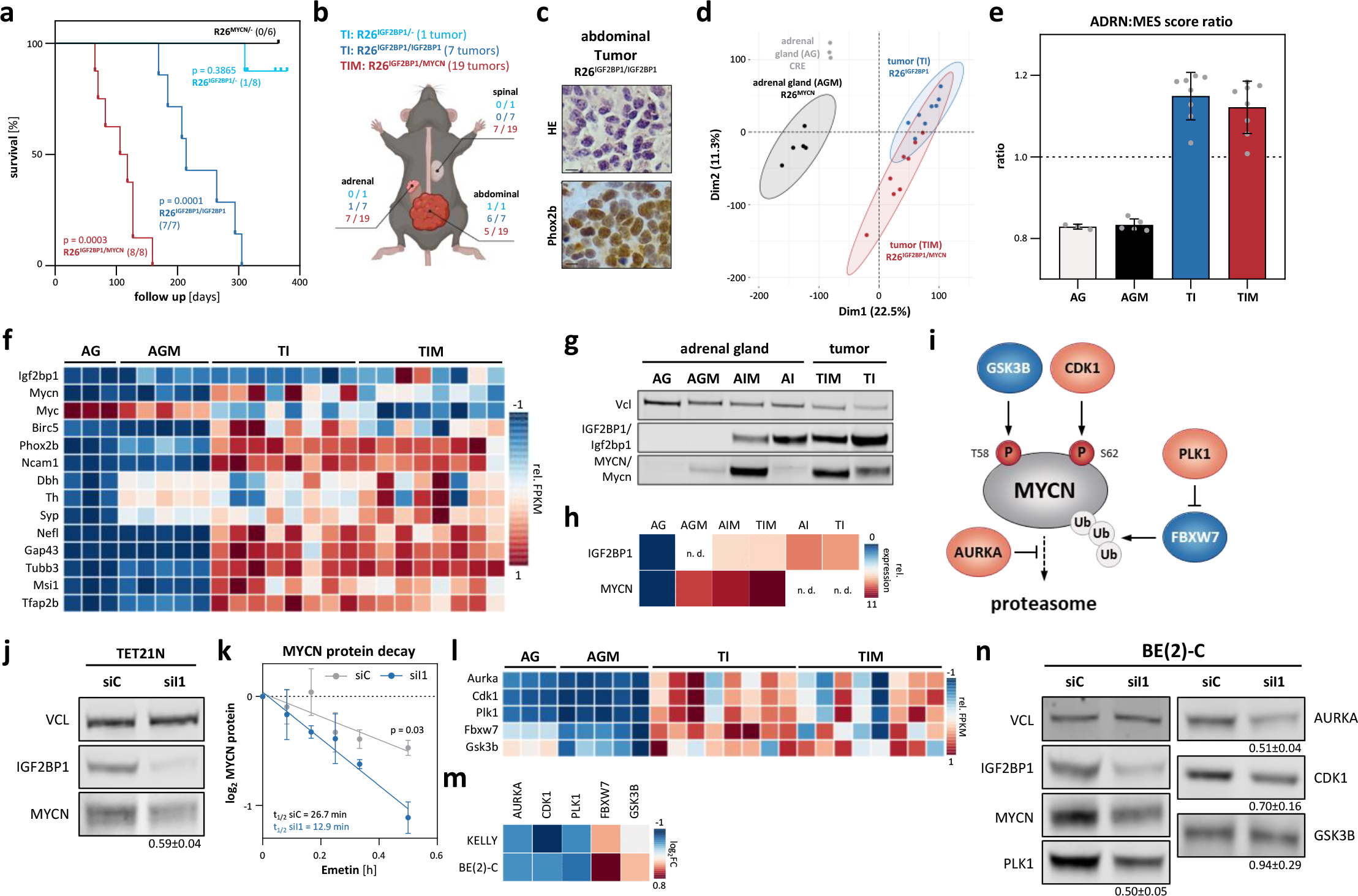
IGF2BP1 induces neuroblastoma, Mycn expression, stabilizes MYCN protein and synergizes with MYCN in transgenic mice. **(a)** Kaplan-Meier survival analysis of heterozygous (cyan, n = 8) and homozygous (blue, n = 7) R26^IGF2BP1^, heterozygous R26^MYCN^ (black, n = 6) and double transgenic R26^IGF2BP1/MYCN^ (red, n = 8) mice. Numbers in brackets indicate tumor bearing mice. **(b)** Scheme of tumor location within mice. **(c)** Representative images of hematoxylin and eosin staining (HE, top) and Phox2b immunohistochemistry (bottom) indicative for neuroblastoma in R26^IGF2BP1/IGF2BP1^ mice (bars: 40 µm). **(d)** PCA of mouse adrenal glands and transgenic tumors. **(e)** Ratio of ADRN to MES signature of mouse adrenal glands and transgenic tumors. **(f)** Heatmap depicting row-scaled FPKM values of indicated murine mRNAs in adrenal glands derived from wildtype (n = 3, AG) or R26^MYCN/-^ (n = 5, AGM) mice and tumors of R26^IGF2BP1^ (n = 8, TI) or R26^IGF2BP1/MYCN^ (n = 8, TIM) mice. **(g)** Western blot analysis confirms IGF2BP1 or MYCN transgene expression in tumors and adrenal glands of representative mice (n = 1). **(h)** RT-qPCR analysis of human IGF2BP1 and MYCN mRNA in indicated mouse tissue (n. d. - not determined). **(i)** Scheme of MYCN protein regulation. **(j)** Western blot (n = 5) analysis of MYCN expression upon IGF2BP1 (siI1) compared to control (siC) knockdown in TET21N cells. **(k)** MYCN protein decay was monitored by Western Blot analysis in control (grey) and IGF2BP1 knockdown (blue) TET21N cells upon indicated time of emetin treatment (n = 3). (l, m) RNA-seq analysis of indicated mRNAs in mouse tissues **(l)** or upon transient IGF2BP1 knockdown in BE(2)-C and KELLY **(m)**. **(n)** Western blot (n = 3) analysis of indicated proteins upon IGF2BP1 (siI1) compared to control (siC) knockdown in BE(2)-C.

### IGF2BP1 induces high-risk neuroblastoma with 2p/17q syntenic chromosomal aberration

The genomic landscape of MYCN/IGF2BP1-induced neuroblastoma was analyzed by sWGS performed on AGM, TI, TIM and murine wildtype adrenal glands. Barely any expanded chromosomal aberrations were observed in AGM, supporting lack of tumor induction (Supplementary Fig. 14a). TIM samples only showed modest (<20 %) gains at Chr 11 and pronounced (up to 50%) gains at Chr 6, whereas barely any deletions were observed (Fig. 7a). In contrast, chromosomal deletions (mainly at Chr 4, 6, 9, 14, and 16) were expanded in TI samples (Fig. 7b). Most strikingly, however, substantial gains (>50- 70 %) were found at Chr 11 (*Igf2bp1*) and Chr 12 (*Mycn*) in TI samples. Chromosomal lift-over analyses (mouse to human) indicated that R26^IGF2BP1^ induces chromosomal gains reminiscent to 2p (*MYCN*) and 17q (*IGF2BP1*) amplification observed in human HRN (Fig. 7c).

**Fig. 7:**
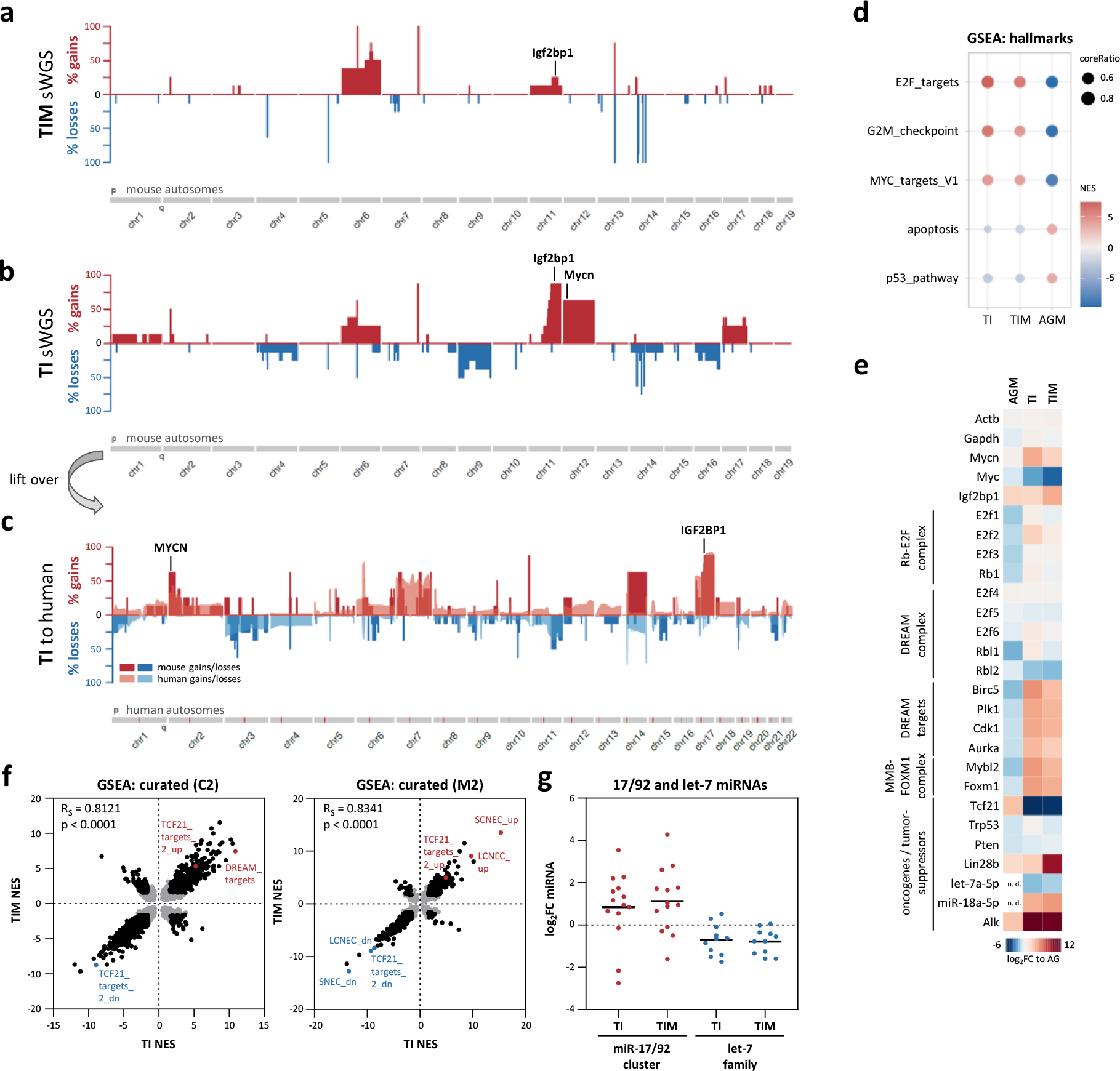
IGF2BP1-induced murine neuroblastoma comprises syntenic chromosomal aberrations and gene expression profiles observed in human high-risk disease. (a-c) Frequency (%) of DNA copy number gains (red) and losses (blue) for murine chromosome 1 to 19 in R26^IGF2BP1/MYCN^ **(a)** or R26^IGF2BP1^ **(b)** tumors compared to wildtype adrenal glands and lift over of R26^IGF2BP1^ regions to the human genome (c, Chr 1-22). Overlay of human neuroblastoma sWGS is depicted in transparent colors **(c)**. **(d)** Selected hallmark gene sets in R26^IGF2BP1^ (TI), R26^IGF2BP1/MYCN^ (TIM) or R26^MYCN^ (AGM) mice based on GSEA. **(e)** RNA-seq analysis of indicated mRNAs presented as log_2_ fold change (log_2_FC) compared to wildtype adrenal glands. **(f)** Correlation of NES values of C2 (left) and M2 (right) gene sets between TI and TIM. Non-significant gene sets are depicted in grey. **(g)** Log_2_FC of miRNAs from the miR-17-92 cluster and let-7 family between R26^IGF2BP1^ or R26^IGF2BP1/MYCN^ tumors and normal adrenal gland tissue. SCNEC/LCNEC - genetic mouse model high-grade small/large-cell neuroendocrine lung carcinoma

GSEA based on RNA-seq data revealed substantial upregulation of pro-proliferative (E2F_TARGETS, G2M_CHECKPOINT) and downregulation of tumor-suppressive (P53_PATHWAY, APOPTOSIS) hallmark gene sets in TI and TIM samples (Fig. 7d; Supplementary Table 3). Consistent with strong elevation of MYCN/Mycn expression, canonical MYC/N-driven gene expression (MYC_TARGETS_V1) was markedly upregulated in tumors. This was in sharp contrast to AGM samples, indicating strong MYCN/Mycn-dependent tumor induction and progression in both, TI and TIM. Intriguingly, strong association of hallmark NES (normalized enrichment score) values was determined in between tumors, but not in comparison to AGM, indicating strong similarity of gene expression in TI and TIM (Supplementary Fig. 14b). This was further evaluated for representative genes and by GSEA of larger curated gene set collections, the human C2 and mouse M2 (Fig. 7e, f; Supplementary Table 3). GSEA in C2 indicated up- and downregulation of gene signatures characteristic for pediatric cancer and MYC/N-driven gene expression (Supplementary Fig. 14c). One of the strongest upregulations in C2 was observed for the FISCHER_DREAM_TARGETS gene set, which partially overlaps with E2F_TARGETS [71]. Despite known cross-talk of MYC/N and E2F-driven gene expression, this supports IGF2BP1-dependent stimulation of RB-E2F-controlled cell cycle (CC) genes and suggests additional roles of IGF2BP1 in activating DREAM-repressed CC genes. Among most downregulated gene sets was CUI_TCF21_TARGETS_2_DN, whereas CUI_TCF21_TARGETS_2_UP was significantly enriched. Accordingly, the expression of TCF21, a crucial developmental transcription factor and strong tumor suppressor [72], was essentially abolished in tumors (Fig. 7e). GSEA in M2 confirmed TCF21-dependency and revealed similarities with high-grade small- (SCNEC) and large-cell (LCNEC) neuroendocrine lung carcinoma mouse models. These where derived by combined deletion of key tumor suppressors (Pten and Trp53) and disturbing E2F-/DREAM-repression of CC genes by deleting Rb1 or Rbl1/Rbl2 (p107/p130) [73]. Only Rbl2 was substantially decreased in tumors, whereas Trp53, Rb1, Rbl1 and Pten mRNA expression remained largely unchanged (Fig. 7e). The downregulation of Rbl2, a key target of the MYC/N-driven miR-17/92 cluster [74], prompted us to test miRNA expression in TI and TIM. Consistent with upregulation of MYCN/Mycn-driven gene synthesis, the pro-tumorigenic miR-17/92 cluster was substantially upregulated in both (Fig. 7e, g). In contrast, the rather tumor suppressive let-7 miRNA family was decreased, supporting modest to strong upregulation of the let-7 miRNA suppressor Lin28b in tumors (Fig. 7e).

## DISCUSSION

Our studies provide the first evidence that IGF2BP1 is a potent and druggable oncogene in neuroblastoma, with target prospects in other MYC/N-driven malignancies. In synergy with MYCN, IGF2BP1 initiates and promotes HRN by transcriptional/post-transcriptional feedforward regulation. MYCN is a versatile transcriptional driver of oncogene expression, most prominently in MNA neuroblastoma [2]. IGF2BP1 is a post-transcriptional enhancer of oncogene expression [21–24], including MYCN and BIRC5, in various cancers with exceeding expression in Chr 17q unbalanced neuroblastoma. MYCN/IGF2BP1 cross-talk results in fostered MYCN-driven transcription and IGF2BP1-directed mRNA stabilization of oncogenic effectors. Many of these, e.g. *NME1*, *PYCR1*, *TK1*, *BIRC5* and *TOP2A*, are located on Chr 17q, are upregulated in unfavorable neuroblastoma and associated with poor patient outcome [12, 18, 75–77]. Strikingly, MYCN/IGF2BP1-driven gene expression leads to increased output of oncogenic factors in a largely miRNA-dependent manner [53, 78, 79]. Most prominently, IGF2BP1 impairs the MYC/N-driven miR-17/92 cluster in down tuning MYC/N expression by negative feedback regulation [53, 74]. MYCN/IGF2BP1 feedback regulation also stimulates LIN28B expression, a strong inducer of neuroblastoma [66]. This promotes oncogenic factors like MYCN, IGF2BP1 and LIN28B by decreasing let-7 miRNA biogenesis and enhancing miRNA antagonizing roles of IGF2BP1 [22, 80]. Thus, deregulated miRNA synthesis and selective impairment of miRNA-directed downregulation of oncogene expression by IGF2BP1 are major mechanisms promoting neuroblastoma initiation and progression. IGF2BP1 furthermore stabilizes MYCN protein, apparently by enhancing E2F/DREAM-regulated genes like AURKA, PLK1 and CDK1 [24, 54, 71]. Collectively, this emphasizes the strong synergy of MYCN and IGF2BP1 in promoting HRN.

MYCN/IGF2BP1 synergy also promotes chromosomal gains at 2p and 17q, but retains strong potential also in non-amplified (2p or 17q) neuroblastoma models, e.g. NBL-S. This relies on the broad impact of MYCN/IGF2BP1-driven gene expression leading to enforced oncogene synthesis, most prominently of Chr 17q genes like *BIRC5*. In transgenic LSL-*IGF2BP1* mouse models, neuroblastoma formation was strictly IGF2BP1 dose-dependent. High incidence was only observed at elevated IGF2BP1 expression in homozygous mice. Apparently, feedforward regulation by MYCN/IGF2BP1 critically relies on IGF2BP1 to stabilize the exceeding demand of transcripts arising by MYCN-driven synthesis. Accordingly, MYCN like MYC [56] promotes IGF2BP1 synthesis culminating in an oncogene storm, which likely contributes to severe genome destabilization. This is evidenced by chromosomal misbalance reminiscent of human neuroblastoma in MYCN- as well as IGF2BP1-driven neuroblastoma mouse models [65, 81]. In support of dose dependence and IGF2BP1-directed MYCN protein stabilization, this is not seen in *R26^MYCN^* model, but striking in R26^IGF2BP1^, the only model with Chr 2p/17q syntenic aberrations. This suggests that super-critical IGF2BP1 levels require a second hit, most likely *MYCN*-amplification, feeding IGF2BP1 synthesis to push feedforward regulation to a level unleashing an oncogene storm. Strong oncogenic synergy of MYCN/IGF2BP1 is evidenced in R26^IGF2BP1/MYCN^ mice, showing reduced tumor latency without or mild chromosomal disturbance. A simple matter of time appears unlikely in view of genomic aberrations observed in other *MYCN*- driven neuroblastoma models within 80 d [65]. Irrespectively, R26^IGF2BP1/MYCN^ and R26^IGF2BP1/IGF2BP1^ neuroblastoma show strikingly similar gene expression, most prominently induction of E2F/DREAM-controlled CC genes. This strongly emphasizes the impact of MYCN/IGF2BP1 synergy in reprogramming gene expression independent of underlying genomic disturbances. IGF2BP1 fosters resistance to apoptosis by DNA-damaging drugs, e.g. YM-155 and doxorubicin [14, 55]. This may also limit spontaneous proliferation-associated apoptosis due to IGF2BP1-dependent elevation of cIAP (BIRC2), survivin (BIRC5), Wip-1 (PPM1D) and TOP2A, which promote DNA-repair and/or serve roles in chromosome stability as well as apoptosis [82–85]. Along these lines, it appears reasonable that IGF2BP1 supports MYC/N-driven proliferation capacity in embryonal neuroblasts, other childhood/youth cancers with strong MYC/N-dependency and aberration at Chr 17q, e.g. rhabdomyosarcoma and medulloblastoma, as well as carcinomas in general.

Consistent with devastating consequences of MYCN/IGF2BP1 synergy, our studies highlight potential therapeutic prospects of IGF2BP1i. Its dose-dependent and broad oncogenic action indicates promising target potential of IGF2BP1 for pharmacological inhibition. Even if IGF2BP1i by small molecule inhibitors disrupting mRNA stabilization remains incomplete, as typically observed, it will have broad impact. IGF2BP1i likely affects many so far non-druggable effectors and directly (e.g. BIRC5) or largely indirectly (e.g. MYC/N) targetable oncogenes. This would be expected to reduce adverse effects and enhance therapeutic efficacy in combined therapies, e.g. shown here with BIRC5i and previously suggested with TOP2Ai [14]. In support of this, IGF2BP1i by BTYNB was proven beneficial in combined treatment with standard-of-care drugs for neuroblastoma treatment like Etoposide or Vincristine [55]. Moreover, BTYNB promotes the efficacy of therapies impairing E2F/DREAM-regulated CDK4/6, e.g. by Palbociclib [24], and indirect inhibition of MYCN by BRDi [58, 59]. Among a variety of BRDi, we demonstrate the best synergy of IGF2BP1i with Mivebresib. Here, we provide the first *in vivo* evidence for such benefits by revealing that BTYNB is a promising lead for developing more potent IGF2BP1i in cancer treatment. In neuroblastoma xenograft studies, BTYNB pre-treatment impaired engraftment and growth. Strikingly, i.t. BTYNB application alone reduced tumor growth without substantial side effects. However, monotherapies with i.p. application of BTYNB on PDX model resulted in less prominent efficacy probably due to high plasma protein binding and potentially poor pharmacological properties.

## Conclusion

We provide the first evidence that IGF2BP1 is a potent and druggable chromosome 17q oncogene. In neuroblastoma, IGF2BP1 synergizes with MYCN by transcriptional/post-transcriptional feed-forward regulation, which unleashes an oncogene storm and genomic instability reminiscent of high-risk diseases. Our studies unravel that impairing IGF2BP1-RNA association by the small molecules BTYNB disrupts this oncogene storm and provides benefits in combined treatment with MYC/N inhibition via BRD inhibitors, most prominently Mivebresib, as well as inhibitors of MYCN/IGF2BP1-driven oncogenes like BIRC5. Future studies need to progress the potency and pharmacological properties of IGF2BP1 inhibitors to effectively disrupt IGF2BP1-dependent oncogene enhancement. BTYNB appears a promising starting point for these endeavors. The here presented mouse models will expedite the evaluation of improved IGF2BP1i and provide valuable resources for identifying additional IGF2BP1-driven oncogenic effectors for combined cancer treatment.

## Abbreviations

3’UTR: 3’ untranslated region; ADRN: adrenergic; AG: adrenal gland; AGM: adrenal gland from LSL-MYCN mouse model; AGO2: argonaute RISC catalytic component 2; AI: adrenal gland from LSL- IGF2BP1 mouse model; AIM: adrenal gland from LSL-IGF2BP1/MYCN mouse model; BIRC5: baculoviral IAP repeat containing 5; BRD: bromodomain-containing protein; BW: body weight; CC: cell cycle; ChIP: chromatin immunoprecipitation; Chr: chromosome; CN: copy number; CRISPR/Cas9: clustered regularly interspaced short palindromic repeats and CRISPR associated protein 9; DBH: dopamine beta-hydroxylase; E2F1: E2F transcription factor 1; EC_50_: half maximal effective concentration; eCLIP: enhanced crosslinking and immunoprecipitation; EEF2: eukaryotic translation elongation factor 2; FACS: fluorescence-activated cell sorting; GAPDH: glyceraldehyde-3-phosphate dehydrogenase; GSEA: gene set enrichment analysis; hESCs: human embryonic stem cells; HIST1: histone cluster 1 H2A family member c; HIST2: histone cluster 2 H3 family member a; HPLC: high performance liquid chromatography; HR: hazard ratio; HRN: high-risk neuroblastoma; i.p.: intra- peritoneal; i.t.: intra-tumoral; iCRE: improved Cre recombinase; IGF2BP1: insulin-like growth factor 2 mRNA-binding protein 1; INSS: international neuroblastoma staging system; KO: knockout; LCNEC: high-grade large-cell neuroendocrine lung carcinoma; m^6^A: N^6^-methyladenosine; MES: mesenchymal; METTL14: methyltransferase 14, N^6^-adenosine-methyltransferase subunit; METTL3: methyltransferase 3, N^6^-adenosine-methyltransferase complex catalytic subunit; miTRAP: miRNA trapping by RNA in vitro affinity purification; MNA: *MYCN*-amplified; MS2-BP: MS2-binding protein; mut: mutated; MYCN: v-myc avian myelocytomatosis viral oncogene neuroblastoma derived homolog; NES: normalized enrichment score; nMNA: non-*MYCN*-amplified; PCA: principal component analysis; PDX: patient-derived xenograft; R26: ROSA26 locus; RIP: RNA immunoprecipitation; RPLP0: ribosomal protein lateral stalk subunit P0; s.c.: subcutaneous; SCNEC: high-grade small-cell neuroendocrine lung carcinoma; sWGS: shallow whole genome sequencing; TI: tumor from LSL-IGF2BP1 mouse model; TIM: tumor from LSL-IGF2BP1/MYCN mouse model; VCL: vinculin; wt: wildtype; ZIP: zero interaction potency

## Declarations

Ethical approval and consent to participate Neuroblastoma tumors were granted after patient consent and ethical approval from the Cologne tumor bank and the Universitätsklinikum Essen, Germany. Animals were handled according to the guidelines of the Martin Luther University. Permission was granted by the state administration office of Saxony-Anhalt (reference number: 42502-2-1381 MLU, 42502-2-1530 MLU, 42502-2-1625 MLU).

## Consent for publication

Not applicable.

## Availability of data and materials

Presented sequencing data have been deposited in NCBI’s Gene Expression Omnibus SuperSeries and are accessible through GEO accession number GSE181582 (https://www.ncbi.nlm.nih.gov/geo/query/acc.cgi?acc=GSE181582). Human DNA- and RNA-seq data are also available via the R2: Genomics Analysis and Visualization Platform (http://r2.amc.nl; datasets: “Tumor Neuroblastoma - Bell”) for interactive use. Corresponding miRNA sequencing data for human neuroblastoma samples is available under GEO accession number GSE155945 or in the R2 database. Processed sequencing data for all analysis (transgenic mice, human neuroblastoma, cell lines) are available in Supplementary Table 9.

## Authorś contributions

S. Hagemann, D.M., M.I.L. and S. Hüttelmaier contributed to project coordination, data analysis and manuscript writing. S. Hagemann and J.L.B. performed experiments and data analysis. D.M. performed bioinformatics analyses. T.F. and S. Hagemann performed animal experiments. J.L.B. and J.H.S. recruited human tumor samples and their accompanying clinical data. N.B. prepared the images for IHC. M.H. performed histological analysis of mouse tumors. E.G. and W.S. provided BTYNB and serum analyses.

## Acknowledgements

We thank Frank Berthold, Barbara Hero and staff at Cologne Neuroblastoma Tumor Bank (Germany) for providing tumor specimens for the here presented studies research. We kindly thank Jacob Haase, Marie-Luise Conrad and Raseswari Turlapati for assistance in tumor preparation. We thank the Institute of Pathology for embedding mouse tissue into paraffin blocks and Beate Heydl for assistance with IHC staining. We thank Jan Koster for implementing the sequencing data into the R2 database (http://r2.amc.nl). We thank Dr. Dennis Gürgen and the EPO company (Berlin, Germany) for performing the treatment experiment in PDX neuroblastoma model. We thank Dr. Knut Krohn (IZFK Leipzig) for assisting with initial sequencing run of human neuroblastoma samples. We thank Dr. Matthias Schmidt for the HPLC analyses.

## Supplementary information

Additional file 1 Supplementary Fig. 1: IGF2BP1 and MYCN upregulation synergize in high-risk neuroblastoma. (a-h) Kaplan-Meier survival analyses by indicated conditions (best cut-off). (i) Spearman correlation of IGF2BP1 and MYCN mRNA expression in neuroblastoma tumors. MNA samples are color-coded in red. (j-m) IGF2BP1 mRNA expression separated by MYCN status (j), Chr 17 balance status (k), INSS stage (l) and survival (m). (n) Correlation analysis of Spearman correlation coefficients determined for IGF2BP1- and MYCN-associated gene expression in primary tumors. (o) Spearman correlation analysis of the logFC of mRNAs after IGF2BP1- or MYCN-knockdown in BE(2)-C. (p) Expression of IGF2BP1 (middle) and MYCN (right) in neuroblastoma upon single cell sequencing. Cell types are indicated in the left panel. (q) Correlation analysis of single cell sequencing data of IGF2BP1 and MYCN (left) or ACTB (right). nMNA - MYCN non-amplified, MNA - MYCN-amplified, bal - balanced Chr 17, unbal - unbalanced Chr 17

Additional file 2 Supplementary Fig. 2: IGF2BP1-driven and BTYNB-druggable MYCN expression is conserved in NBL-S and KELLY neuroblastoma cells. (a) IGF2BP1-CLIP hits in the MYCN mRNA derived from two studies in hESC. (b) Western blot and RT-qPCR analyses of MYCN expression upon I1-KO in NBL-S and KELLY (n = 3). (c) IGF2BP1-RIP analyses in NBL-S and KELLY (n = 3). (d) BTYNB- response curves in NBL-S and KELLY (n = 3). (e) Western blot analysis of MYCN expression upon treatment of NBL-S and KELLY with BTYNB (n = 3). (f-h) Relative activity of MYCN 3’UTR luciferase reporter in NBL-S and KELLY determined between I1-KO and control cells (f), mutant and wildtype MYCN 3’UTR (g) or BTYNB- and DMSO-treated cells (h; all n = 3).

Additional file 3 Supplementary Fig. 3: Validation of IGF2BP1 and AGO2 immunoprecipitation. (a-c) Isolation of indicated proteins (IGF2BP1, AGO2) in RIP studies from parental BE(2)-C, NBL-S and KELLY (a), DMSO- and BTYNB-treated BE(2)-C (b) or parental (Ctrl) and I1-KO (c) cells was analyzed by Western blotting (a, n = 5; b and c, n = 3). BC - bead control

Additional file 4 Supplementary Fig. 4: Regulation of MYCN by IGF2BP1 is largely m^6^A-independent. (a) N^6^-Methyladenosine (m^6^A) modification profile of MYCN mRNA. (b, c) Western blot (b; BE(2)-C, n = 3; KELLY, n = 4) and RT-qPCR (c; n = 3) analyses of MYCN expression upon co-depletion of the key mRNA m^6^A-methyltransferase complex METTL3/14 (siM3/14) compared to control knockdown (siC). (d) RNA-seq data of E2F1 expression in BE(2)-C and KELLY upon transient control (siC) and IGF2BP1 (siI1) knockdown (n = 3). B - BE(2)-C, K - KELLY

Additional file 5 Supplementary Fig. 5: Oncogenic potential of IGF2BP1 is conserved in nMNA neuroblastoma cell and xenograft model. (a, b) The viability and caspase3/7 activity of parental (Ctrl) and I1-KO NBL-S was analyzed in spheroid growth (a; n = 3) and anoikis resistance studies (b; n = 3; bars a, 200 µm; b, 400 µm). (c-e) Tumor growth (n = 6) of Ctrl and I1-KO NBL-S s.c. xenografts was monitored by non-invasive infrared imaging (c), tumor-free survival (d), and final tumor mass (e). (f) RT-qPCR analysis of MYCN mRNA levels in excised parental xenograft tumors (n = 6) and residual non-palpable I1-KO tumors (n = 5).

Additional file 6 Supplementary Fig. 6: BTYNB is stable, has minimal effect on mouse body weight, but high serum protein binding capacity. (a, b) Body weight analysis of (a, n = 6; b, n = 9) mice with s.c. PDX (a) or BE(2)-C xenograft tumors (b) treated i.p. (a) or i.t. (b) with DMSO (grey) or BTYNB (blue). Treatment cycles are indicated by dashed lines below the x-axis. (c) Waterfall plot showing the percentage difference in tumor size from treatment day 0 to day 8 in BE(2)-C xenograft mice treated i.t. with DMSO (beige) or BTYNB (blue; n = 9). (d) Final relative tumor volume of s.c. BE(2)-C xenografts tumors treated with DMSO or BTYNB (n = 9). (e-h) Superimposed HPLC chromatograms for BTYNB stability in DMEM (e), under acidic conditions (f) and after incubation with FBS (g) or HSA (h) at different time points.

Additional file 7 Supplementary Fig. 7: MYCN-driven and BRDi-directed inhibition of IGF2BP1 synthesis is conserved in neuroblastoma cell models. (a) Western blot (n = 3) and RT-qPCR (n = 6) analysis of IGF2BP1 expression upon MYCN (siMN) compared to control knockdown (siC) in KELLY. (b) BRD inhibitor response curve in BE(2)-C, KELLY and NBL-S (n = 4). (c) Western blot (KELLY, n = 3; NBL- S, n = 4) and RT-qPCR (n = 3) analysis of MYCN and IGF2BP1 expression after treatment of KELLY or NBL-S with indicated BRD inhibitors. (d) BE(2)-C cell viability upon treatment with optimal synergy concentrations (750 nM BTYNB, 10 nM Mivebresib) alone or in combination. BTY - BTYNB, Mive - Mivebresib. (e, f) Relief plot showing the ZIP synergy for combined treatment of BTYNB and Mivebresib in KELLY (e) and NBL-S (f; n = 3).

Additional file 8 Supplementary Fig. 8: MYCN/IGF2BP1-driven upregulation of BIRC5 is conserved in neuroblastoma models and primary human disease. (a) Heatmap representing the identification of potential IGF2BP1 and MYCN downstream targets. Neuroblastoma and common essential genes located on Chr 17q were analyzed as described (for details refer to the online method section). Data of each row were scaled to range 0 and 1 (ChIP and CLIP) or -1 and 1 (all other). (b) Kaplan-Meier survival analysis of human neuroblastoma by BIRC5 expression (median cut-off). (c, d) Spearman correlation analyses of BIRC5 with MYCN (c) or IGF2BP1 (d) mRNA expression in human neuroblastoma tumor samples. (e-g) Expression of BIRC5 in neuroblastoma separated by MYCN status (e), Chr 17 balance status (f) and INSS tumor stage (g). (h) Western blot analysis of BIRC5 expression upon transient MYCN (siMN) or IGF2BP1 (siI1) compared to control (siC) knockdown in KELLY (n = 5). (i) BIRC5 3’UTR reporter activity in I1-KO versus Ctrl and BTYNB- versus DMSO-treated KELLY (n = 3). (j) Relief plot showing the ZIP synergy for combined treatment of BTYNB and YM-155 in BE(2)-C (n = 3). nMNA - MYCN non-amplified, MNA - MYCN-amplified, bal - balanced Chr 17, unbal - unbalanced Chr 17, TS - tumor suppressor

Additional file 9 Supplementary Fig. 9: Expression of transgenic tumors is largely location-independent. (a) Schematic overview of the LSL-transgene expression of IGF2BP1 and iRFP or MYCN and luciferase respectively from the Rosa26 locus. Location of primers used for genotyping (A1, A2, A3, I1, I2, M1, M2) are indicated. Splice acceptor site (SA), neomycin resistance (5’del NeoR), the synthetic promoter (CAG), transcriptional stop cassette made of the human growth hormone polyadenylation signal (hGHpA), transgene open reading frame (blue), internal ribosomal entry site (IRES), iRFP or luciferase open reading frame (red), polyadenylation signal (pA). (b) Representative genotyping PCR validating the transgene knockin (top, primers A2 and A3) and the wildtype Rosa26 locus (bottom, primers A1 and A3) in heterozygous (+/-), homozygous (+/+) and wildtype (wt) mice. (c) PCA of all sequenced adrenal glands and tumors split by their location. AG - wildtype adrenal gland, AGM - adrenal gland from R26^MYCN^, TI - tumor from R26^IGF2BP1^, TIM - tumor from R26^IGF2BP1/MYCN^

Additional file 10 Supplementary Fig. 10: Summary and validation of analyzed R26^IGF2BP1^ tumors. (a) For each analyzed LSL-IGF2BP1 mouse with obvious signs of tumor burden, expression of IGF2BP1/Igf2bp1, Mycn and Birc5 was validated by Western blotting (left panels, n = 1). Time until termination criteria is reached are indicated in brackets. Mouse 1-6 and 8 had abdominal tumors, whereas mouse 7 had an adrenal gland tumor. HE staining was performed on excised tumors and representative images are shown in right panels (bars 200 µm). (b) Genotyping PCR performed as in Supplementary Figure 10 validated IGF2BP1 transgene (LSL-IGF2BP1), Rosa26 wildtype locus (Rosa26 wt) and Dbh-iCRE. K - kidney, AG - adrenal gland, T - tumor, AG T - adrenal gland tumor

Additional file 11 Supplementary Fig. 11: Summary and validation of analyzed R26^IGF2BP1/MYCN^ tumors. (a) For each analyzed LSL-IGF2BP1/MYCN mouse with obvious signs of tumor burden, expression of IGF2BP1/Igf2bp1, MYCN/Mycn and Birc5 was validated by Western blotting (n = 1). Time until termination criteria is reached are indicated in brackets. (b) Genotyping PCR performed as in Supplementary Figure 10 validated IGF2BP1 and MYCN transgene (LSL-IGF2BP1, LSL-MYCN), absence of Rosa26 wildtype locus (Rosa26 wt) and Dbh-iCRE. K - kidney, AG - adrenal gland, T - tumor, AG T - adrenal gland tumor, T S - tumor along spine, C - control wildtype mouse

Additional file 12 Supplementary Fig. 12: Summary and validation of analyzed R26^MYCN^ adrenal glands. (a) For each analyzed LSL-MYCN mouse expression of Igf2bp1, MYCN/Mycn and Birc5 was validated by Western blotting (n = 1). Each mice were culled after 365 days. As control (C) served an adrenal gland from a R26^IGF2BP1/MYCN^ mouse. (b) Genotyping PCR validated MYCN transgene (LSL-MYCN), Rosa26 wildtype locus (Rosa26 wt) and Dbh-iCRE.

Additional file 13 Supplementary Fig. 13: IGF2BP1 influences MYCN protein turnover. (a) Western blot (n = 3) analysis of MYCN protein decay upon transient IGF2BP1 (siIGF2BP1) or control (siC) knockdown in TET21N. Treatment time with emetin in minutes are indicated. (b) Western blot (n = 3) analysis of indicated phosphoproteins upon IGF2BP1 (siI1) compared to control (siC) knockdown in BE(2)-C. Relative changes of phosphorylation signals to total protein level are indicated.

Additional file 14 Supplementary Fig. 14: Genomic aberration are nearly absent in R26^MYCN^ adrenal glands. (a) Frequency (%) of DNA copy number gains (red) and losses (blue) for murine chromosome 1 to 19 in R26^MYCN^ compared to wildtype adrenal glands. (b) Correlation of NES values of hallmark gene sets between TI and TIM (left) as well as tumors and AGM (middle, right). Non-significant gene sets are depicted in grey. (c) Selected gene sets from curated human (C2) and mouse (M2) collection in R26^IGF2BP1^ (TI), R26^IGF2BP1/MYCN^ (TIM) or R26^MYCN^ (AGM) mice based on GSEA.

Additional file 15 Supplementary Table 1: ADRN/MES neuroblastoma signature.

Additional file 16 Supplementary Table 2: MNA neuroblastoma essential genes and respective dependency score.

Additional file 17 Supplementary Table 3: Gene set enrichment analyses of transgenic mouse tissue/tumors.

Additional file 18 Supplementary Table 4: IGF2BP1 CLIP score for protein-coding genes.

Additional file 19 Supplementary Table 5: MYCN ChIP data for protein-coding genes.

Additional file 20 Supplementary Table 6: Antibodies, primers, plasmids and siRNAs used in this study.

Additional file 21 Supplementary Table 7: Overview of transgenic mice.

Additional file 22 Supplementary Table 8: Overview of the human neuroblastoma cohort.

Additional file 23 Supplementary Table 9: Processed RNA-seq data of human and mouse samples.

