## Supplementary Figures for "IGF2BP1 induces high-risk neuroblastoma and forms a druggable feedforward loop with MYCN promoting 17q oncogene expression"

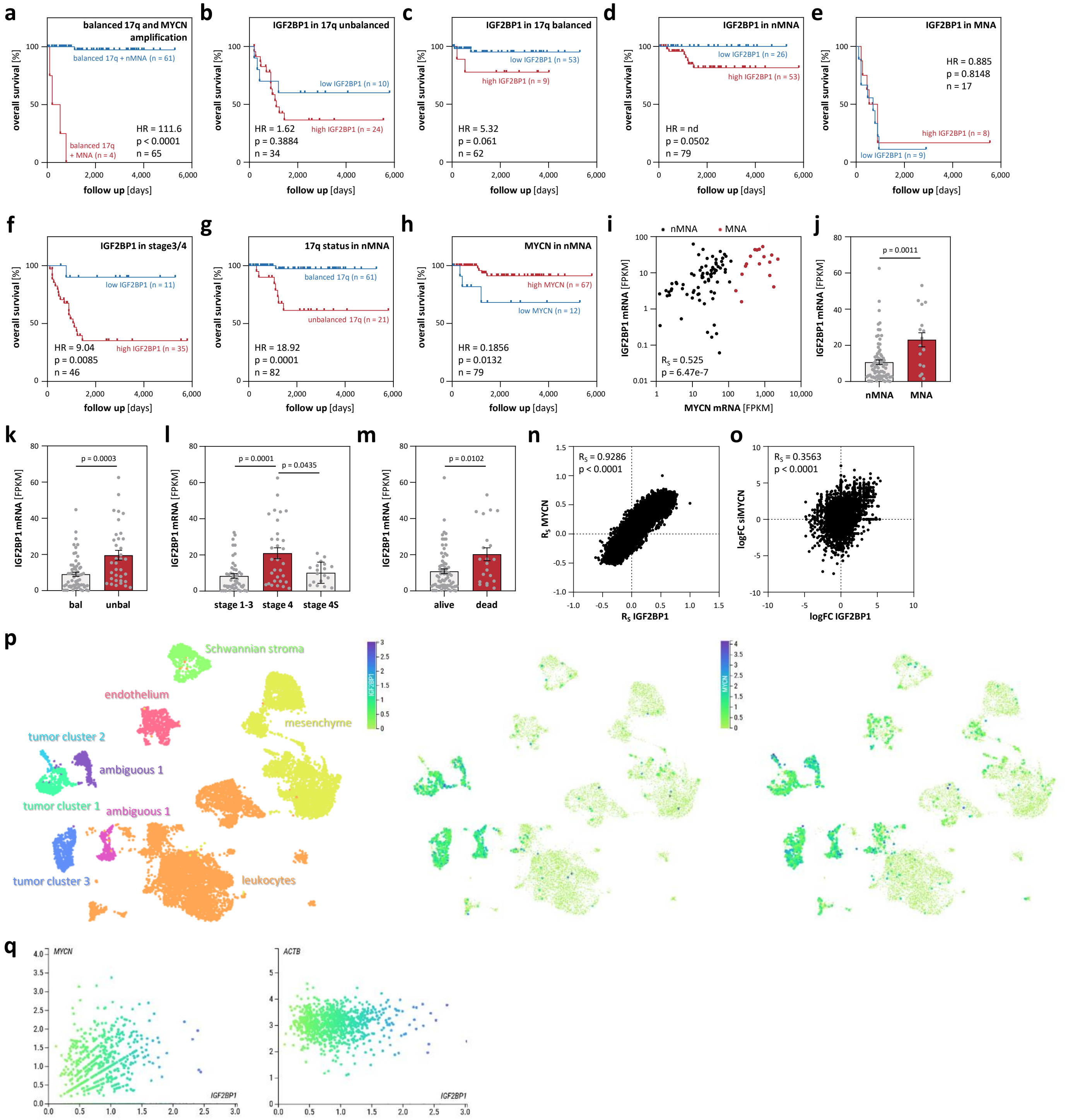
a
balanced 17q and MYCN amplification
balanced 17q + nMNA (n = 61)
balanced 17q + MNA (n = 4)
HR = 111.6
p < 0.0001
n = 65
b
IGF2BP1 in 17q unbalanced
low IGF2BP1 (n = 10)
high IGF2BP1 (n = 24)
HR = 1.62
p = 0.3884
n = 34
c
IGF2BP1 in 17q balanced
low IGF2BP1 (n = 53)
high IGF2BP1 (n = 9)
HR = 5.32
p = 0.061
n = 62
d
IGF2BP1 in nMNA
low IGF2BP1 (n = 26)
high IGF2BP1 (n = 53)
HR = nd
p = 0.0502
n = 79
e
IGF2BP1 in MNA
HR = 0.885
p = 0.8148
n = 17
high IGF2BP1 (n = 8)
low IGF2BP1 (n = 9)
overall survival [%]
follow up [days]
f
IGF2BP1 in stage3/4
low IGF2BP1 (n = 11)
high IGF2BP1 (n = 35)
HR = 9.04
p = 0.0085
n = 46
g
17q status in nMNA
balanced 17q (n = 61)
unbalanced 17q (n = 21)
HR = 18.92
p = 0.0001
n = 82
h
MYCN in nMNA
high MYCN (n = 67)
low MYCN (n = 12)
HR = 0.1856
p = 0.0132
n = 79
i
nMNA
MNA
IGF2BP1 mRNA [FPKM]
MYCN mRNA [FPKM]
RS = 0.525
p = 6.47e-7
j
p = 0.0011
nMNA
MNA
k
p = 0.0003
bal
unbal
l
p = 0.0001
p = 0.0435
stage 1-3
stage 4
stage 4S
m
p = 0.0102
alive
dead
n
RS = 0.9286
p < 0.0001
RS MYCN
RS IGF2BP1
o
RS = 0.3563
p < 0.0001
logFC siMYCN
logFC IGF2BP1
p
Schwannian stroma
endothelium
mesenchyme
tumor cluster 2
ambiguous 1
tumor cluster 1
ambiguous 1
tumor cluster 3
leukocytes
IGF2BP1
MYCN
q
MYCN
IGF2BP1
ACTB
IGF2BP1

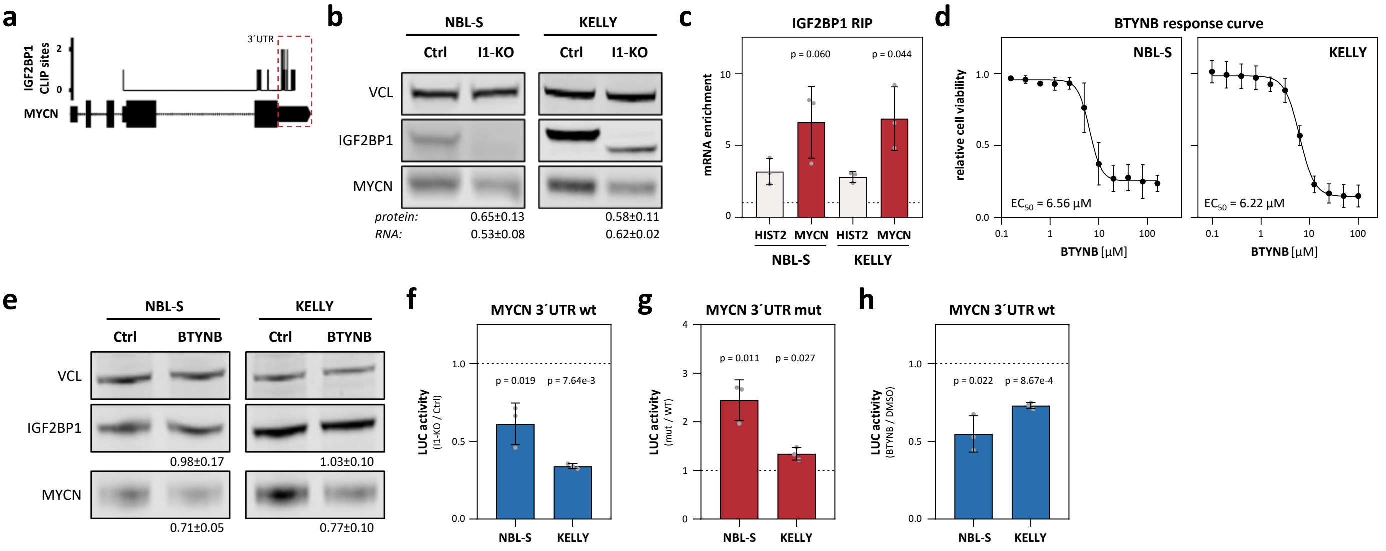
a
IGF2BP1 CLIP sites
3´UTR
MYCN
b
NBL-S
KELLY
Ctrl
I1-KO
VCL
IGF2BP1
MYCN
protein:
0.65±0.13
0.58±0.11
RNA:
0.53±0.08
0.62±0.02
c
IGF2BP1 RIP
mRNA enrichment
p = 0.060
p = 0.044
HIST2
MYCN
d
BTYNB response curve
relative cell viability
EC50 = 6.56 µM
EC50 = 6.22 µM
BTYNB [µM]
e
BTYNB
0.98±0.17
1.03±0.10
0.71±0.05
0.77±0.10
f
MYCN 3´UTR wt
LUC activity
(I1-KO / Ctrl)
p = 0.019
p = 7.64e-3
g
MYCN 3´UTR mut
(mut / WT)
p = 0.011
p = 0.027
h
(BTYNB / DMSO)
p = 0.022
p = 8.67e-4

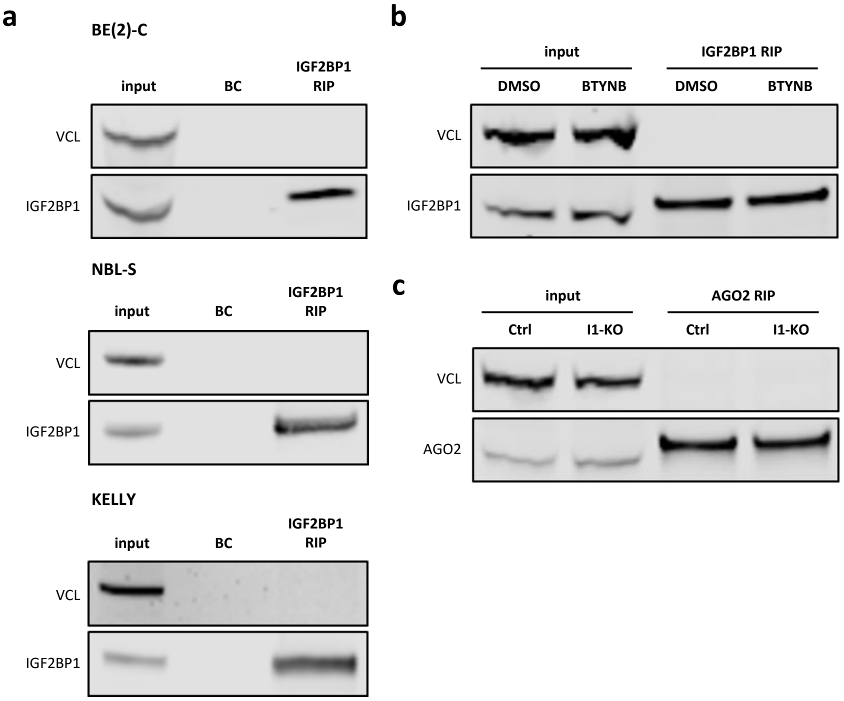
a
BE(2)-C
input
BC
IGF2BP1
RIP
VCL
IGF2BP1
NBL-S
input
BC
IGF2BP1
RIP
VCL
IGF2BP1
KELLY
input
BC
IGF2BP1
RIP
VCL
IGF2BP1
b
input
IGF2BP1 RIP
DMSO
BTYNB
DMSO
BTYNB
VCL
IGF2BP1
c
input
AGO2 RIP
Ctrl
I1-KO
Ctrl
I1-KO
VCL
AGO2

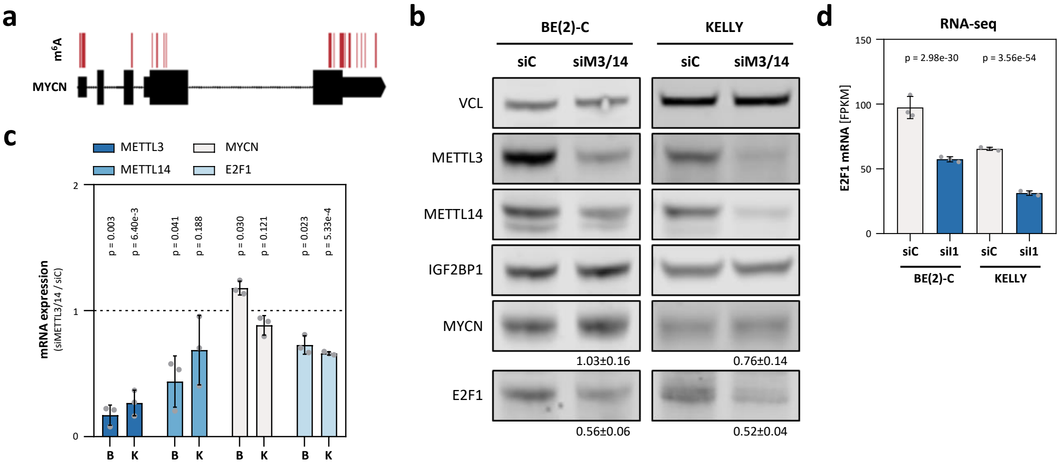

a
m6A
MYCN
b
BE(2)-C
KELLY
siC
siM3/14
siC
siM3/14
VCL
METTL3
METTL14
IGF2BP1
MYCN
1.03±0.16
0.76±0.14
E2F1
0.56±0.06
0.52±0.04
c
METTL3
METTL14
MYCN
E2F1
mRNA expression
(siMETTL3/14 / siC)
p = 0.003
p = 6.40e-3
p = 0.041
p = 0.188
p = 0.030
p = 0.121
p = 0.023
p = 5.33e-4
B
K
B
K
B
K
B
K
d
RNA-seq
p = 2.98e-30
p = 3.56e-54
E2F1 mRNA [FPKM]
siC
siI1
siC
siI1
BE(2)-C
KELLY

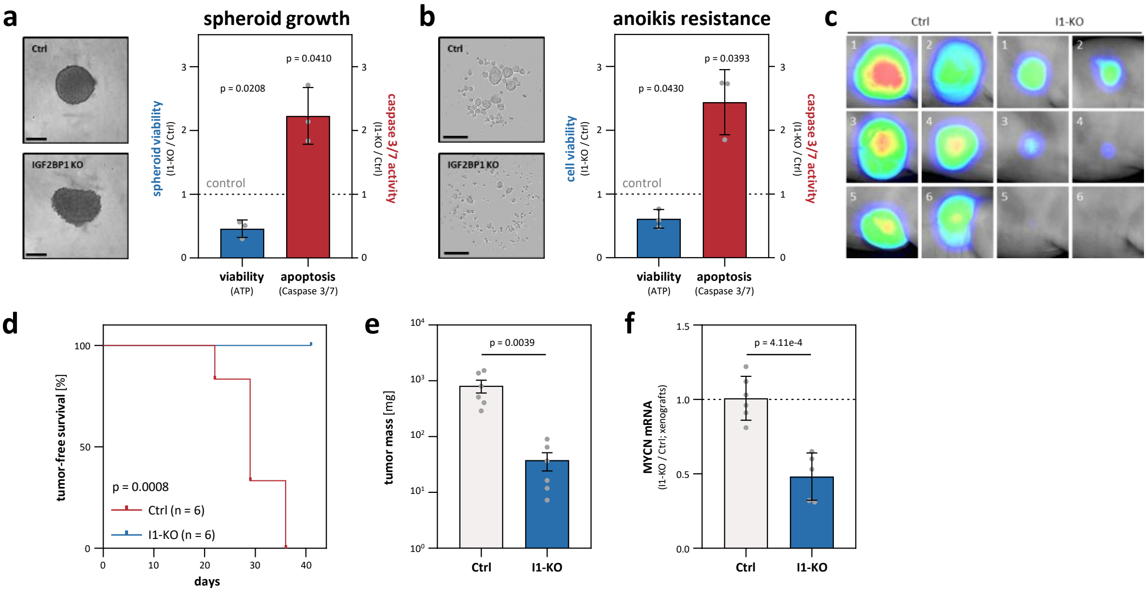

a
spheroid growth
Ctrl
IGF2BP1 KO
spheroid viability
(I1-KO / Ctrl)
caspase 3/7 activity
(I1-KO / Ctrl)
p = 0.0208
p = 0.0410
control
viability
(ATP)
apoptosis
(Caspase 3/7)
b
anoikis resistance
Ctrl
IGF2BP1 KO
cell viability
(I1-KO / Ctrl)
caspase 3/7 activity
(I1-KO / Ctrl)
p = 0.0430
p = 0.0393
control
viability
(ATP)
apoptosis
(Caspase 3/7)
c
Ctrl
I1-KO
d
tumor-free survival [%]
days
p = 0.0008
Ctrl (n = 6)
I1-KO (n = 6)
e
tumor mass [mg]
p = 0.0039
Ctrl
I1-KO
f
MYCN mRNA
(I1-KO / Ctrl; xenografts)
p = 4.11e-4
Ctrl
I1-KO

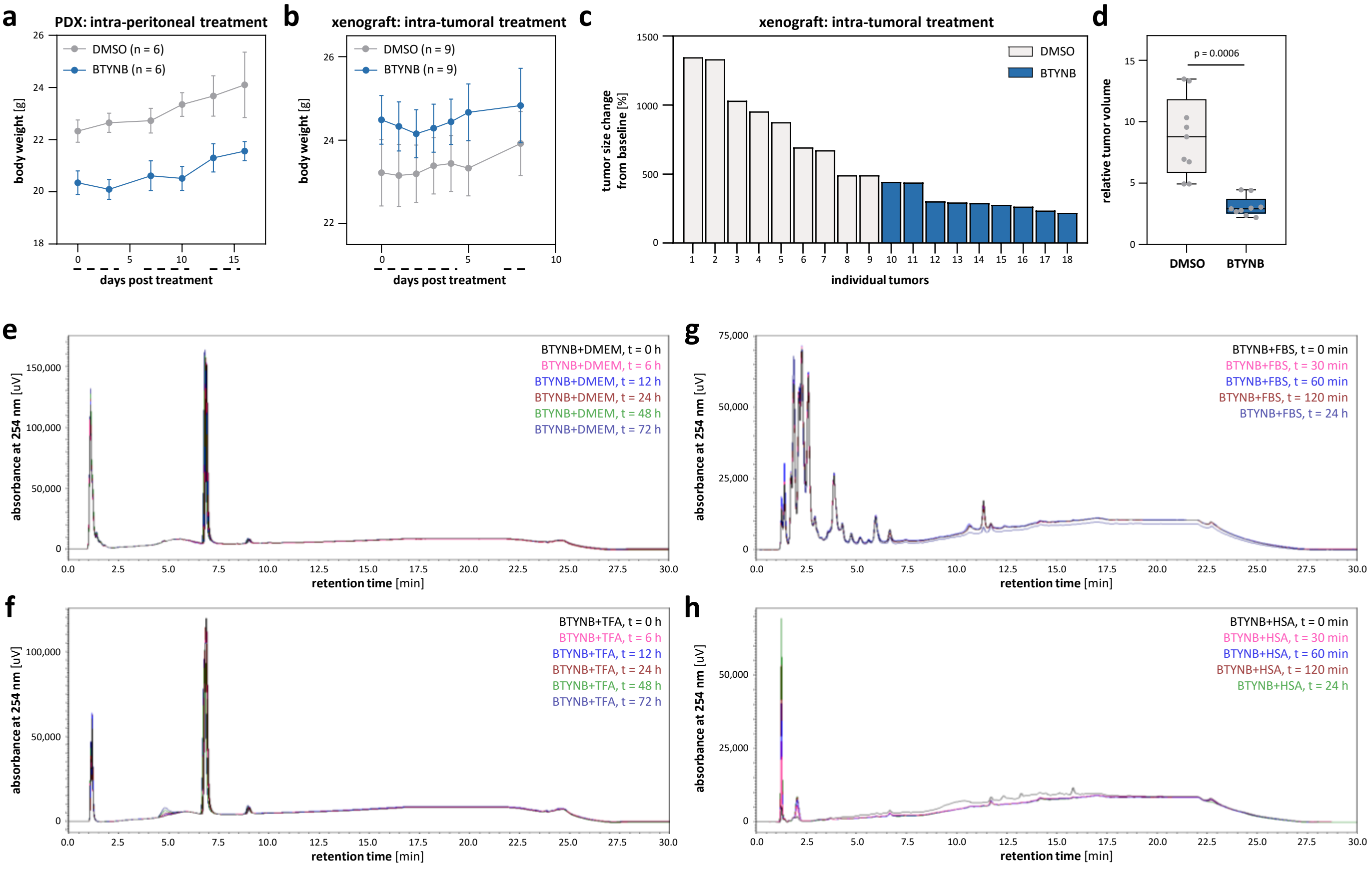

a
PDX: intra-peritoneal treatment
DMSO (n = 6)
BTYNB (n = 6)
body weight [g]
days post treatment
b
xenograft: intra-tumoral treatment
DMSO (n = 9)
BTYNB (n = 9)
body weight [g]
days post treatment
c
xenograft: intra-tumoral treatment
DMSO
BTYNB
tumor size change from baseline [%]
individual tumors
d
p = 0.0006
relative tumor volume
DMSO
BTYNB
e
BTYNB+DMEM, t = 0 h
BTYNB+DMEM, t = 6 h
BTYNB+DMEM, t = 12 h
BTYNB+DMEM, t = 24 h
BTYNB+DMEM, t = 48 h
BTYNB+DMEM, t = 72 h
absorbance at 254 nm [uV]
retention time [min]
f
BTYNB+TFA, t = 0 h
BTYNB+TFA, t = 6 h
BTYNB+TFA, t = 12 h
BTYNB+TFA, t = 24 h
BTYNB+TFA, t = 48 h
BTYNB+TFA, t = 72 h
absorbance at 254 nm [uV]
retention time [min]
g
BTYNB+FBS, t = 0 min
BTYNB+FBS, t = 30 min
BTYNB+FBS, t = 60 min
BTYNB+FBS, t = 120 min
BTYNB+FBS, t = 24 h
absorbance at 254 nm [uV]
retention time [min]
h
BTYNB+HSA, t = 0 min
BTYNB+HSA, t = 30 min
BTYNB+HSA, t = 60 min
BTYNB+HSA, t = 120 min
BTYNB+HSA, t = 24 h
absorbance at 254 nm [uV]
retention time [min]

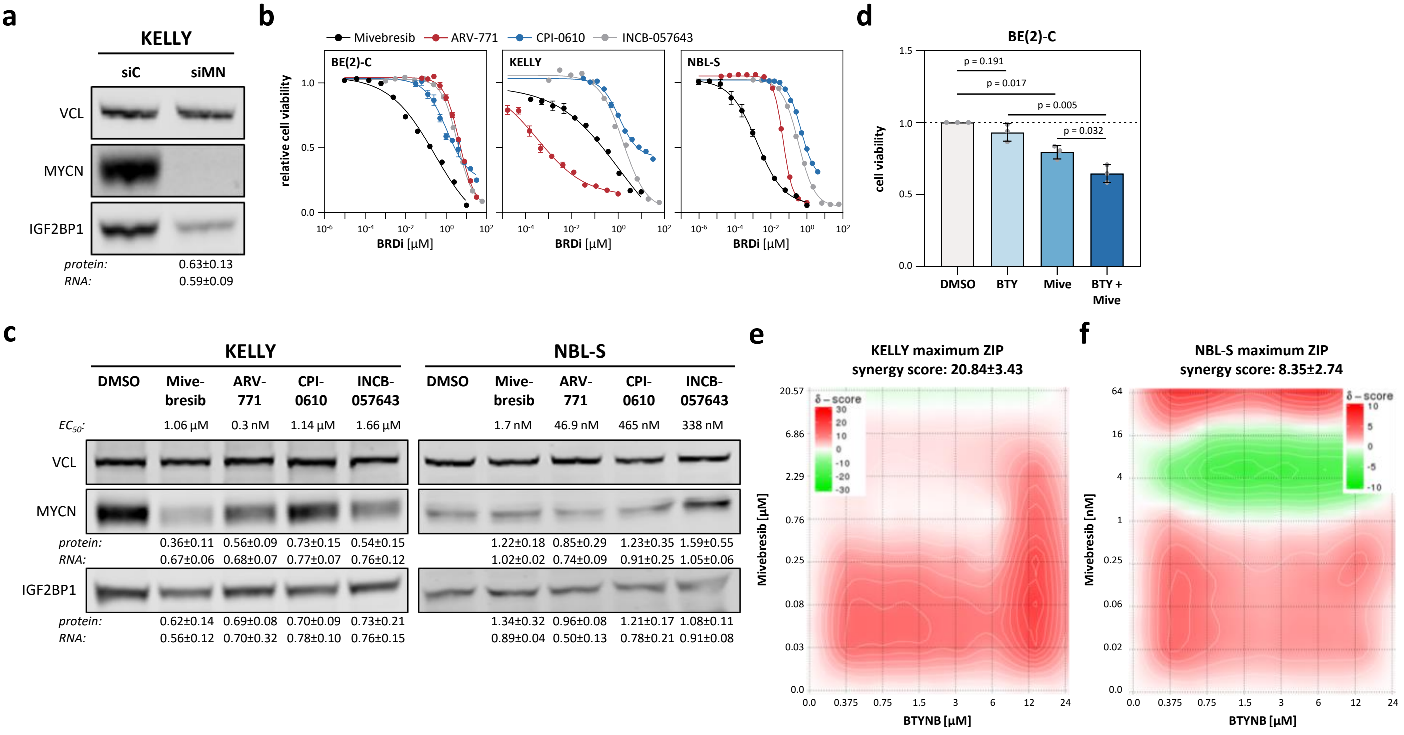
a
KELLY
siC
siMN
VCL
MYCN
IGF2BP1
protein: 0.63±0.13
RNA: 0.59±0.09
b
Mivebresib
ARV-771
CPI-0610
INCB-057643
BE(2)-C
KELLY
NBL-S
relative cell viability
BRDi [µM]
d
BE(2)-C
cell viability
p = 0.191
p = 0.017
p = 0.005
p = 0.032
DMSO
BTY
Mive
BTY + Mive
c
KELLY
NBL-S
DMSO
Mive-bresib
ARV-771
CPI-0610
INCB-057643
EC50: 1.06 µM 0.3 nM 1.14 µM 1.66 µM
1.7 nM 46.9 nM 465 nM 338 nM
VCL
MYCN
protein: 0.36±0.11 0.56±0.09 0.73±0.15 0.54±0.15
RNA: 0.67±0.06 0.68±0.07 0.77±0.07 0.76±0.12
1.22±0.18 0.85±0.29 1.23±0.35 1.59±0.55
1.02±0.02 0.74±0.09 0.91±0.25 1.05±0.06
IGF2BP1
protein: 0.62±0.14 0.69±0.08 0.70±0.09 0.73±0.21
RNA: 0.56±0.12 0.70±0.32 0.78±0.10 0.76±0.15
1.34±0.32 0.96±0.08 1.21±0.17 1.08±0.11
0.89±0.04 0.50±0.13 0.78±0.21 0.91±0.08
e
KELLY maximum ZIP synergy score: 20.84±3.43
δ – score
Mivebresib [µM]
BTYNB [µM]
f
NBL-S maximum ZIP synergy score: 8.35±2.74
δ – score
Mivebresib [nM]
BTYNB [µM]

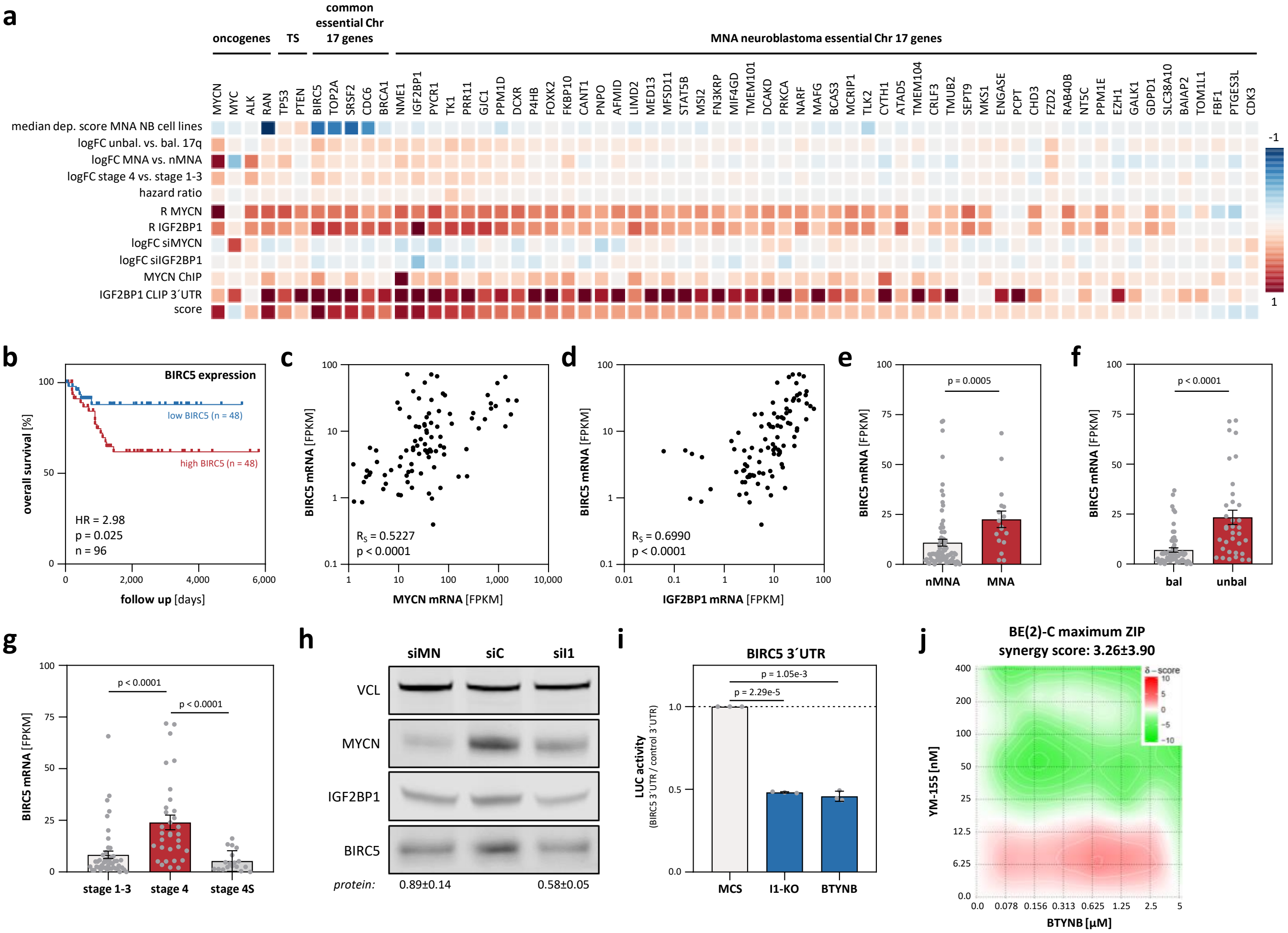

a
oncogenes
TS
common essential Chr 17 genes
MNA neuroblastoma essential Chr 17 genes
median dep. score MNA NB cell lines
logFC unbal. vs. bal. 17q
logFC MNA vs. nMNA
logFC stage 4 vs. stage 1-3
hazard ratio
R MYCN
R IGF2BP1
logFC siMYCN
logFC siIGF2BP1
MYCN ChIP
IGF2BP1 CLIP 3´UTR
score
b
BIRC5 expression
low BIRC5 (n = 48)
high BIRC5 (n = 48)
HR = 2.98
p = 0.025
n = 96
overall survival [%]
follow up [days]
c
$R_S$ = 0.5227
p < 0.0001
BIRC5 mRNA [FPKM]
MYCN mRNA [FPKM]
d
$R_S$ = 0.6990
p < 0.0001
IGF2BP1 mRNA [FPKM]
e
p = 0.0005
nMNA
MNA
f
p < 0.0001
bal
unbal
g
p < 0.0001
p < 0.0001
stage 1-3
stage 4
stage 4S
h
siMN
siC
siI1
VCL
MYCN
IGF2BP1
BIRC5
protein:
0.89±0.14
0.58±0.05
i
BIRC5 3´UTR
p = 1.05e-3
p = 2.29e-5
LUC activity
(BIRC5 3´UTR / control 3´UTR)
MCS
I1-KO
BTYNB
j
BE(2)-C maximum ZIP
synergy score: 3.26±3.90
YM-155 [nM]
BTYNB [µM]
δ - score

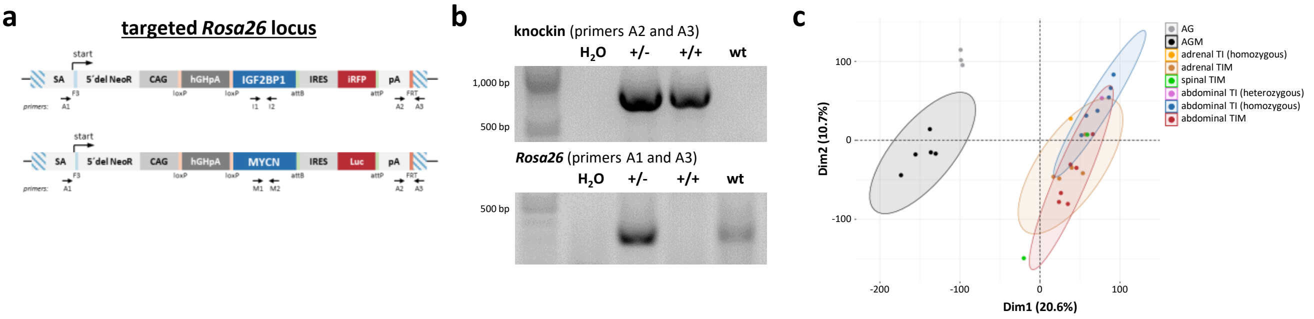

a
targeted Rosa26 locus
start
SA
5´del NeoR
CAG
hGHpA
IGF2BP1
IRES
iRFP
pA
F3
loxP
loxP
attB
attP
FRT
primers:
A1
I1
I2
A2
A3
MYCN
Luc
M1
M2
b
knockin (primers A2 and A3)
H2O
+/-
+/+
wt
1,000 bp
500 bp
Rosa26 (primers A1 and A3)
c
AG
AGM
adrenal TI (homozygous)
adrenal TIM
spinal TIM
abdominal TI (heterozygous)
abdominal TI (homozygous)
abdominal TIM
Dim2 (10.7%)
Dim1 (20.6%)
100
0
-100
-200

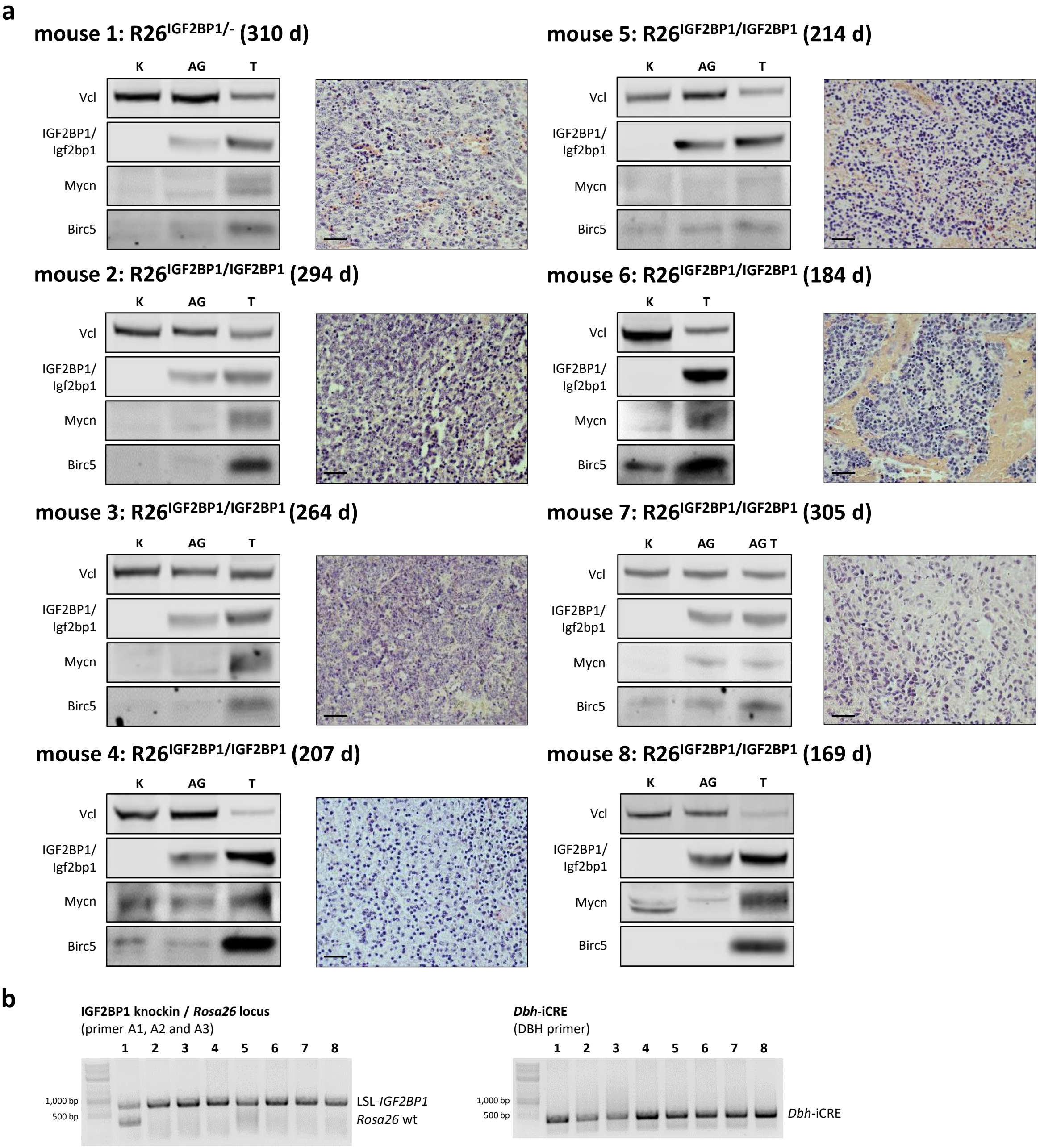
a
mouse 1: R26IGF2BP1/- (310 d)
mouse 2: R26IGF2BP1/IGF2BP1 (294 d)
mouse 3: R26IGF2BP1/IGF2BP1 (264 d)
mouse 4: R26IGF2BP1/IGF2BP1 (207 d)
mouse 5: R26IGF2BP1/IGF2BP1 (214 d)
mouse 6: R26IGF2BP1/IGF2BP1 (184 d)
mouse 7: R26IGF2BP1/IGF2BP1 (305 d)
mouse 8: R26IGF2BP1/IGF2BP1 (169 d)
K
AG
T
AG T
Vcl
IGF2BP1/
Igf2bp1
Mycn
Birc5
b
IGF2BP1 knockin / Rosa26 locus
(primer A1, A2 and A3)
1 2 3 4 5 6 7 8
1,000 bp
500 bp
LSL-IGF2BP1
Rosa26 wt
Dbh-iCRE
(DBH primer)
Dbh-iCRE

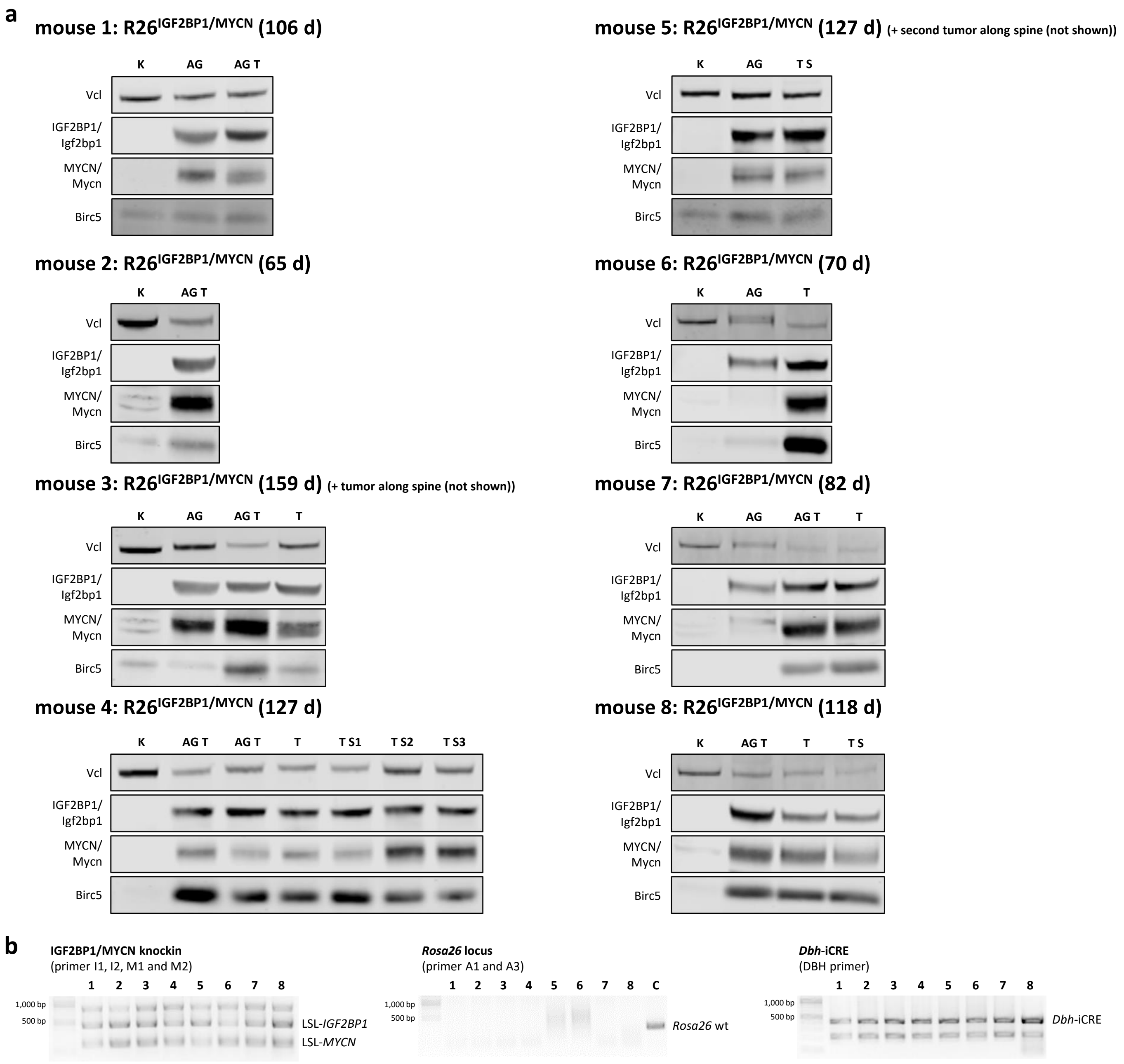
a
mouse 1: R26IGF2BP1/MYCN (106 d)
K
AG
AG T
Vcl
IGF2BP1/
Igf2bp1
MYCN/
Mycn
Birc5
mouse 2: R26IGF2BP1/MYCN (65 d)
mouse 3: R26IGF2BP1/MYCN (159 d) (+ tumor along spine (not shown))
T
mouse 4: R26IGF2BP1/MYCN (127 d)
T S1
T S2
T S3
mouse 5: R26IGF2BP1/MYCN (127 d) (+ second tumor along spine (not shown))
T S
mouse 6: R26IGF2BP1/MYCN (70 d)
mouse 7: R26IGF2BP1/MYCN (82 d)
mouse 8: R26IGF2BP1/MYCN (118 d)
b
IGF2BP1/MYCN knockin
(primer I1, I2, M1 and M2)
1,000 bp
500 bp
1 2 3 4 5 6 7 8
LSL-IGF2BP1
LSL-MYCN
Rosa26 locus
(primer A1 and A3)
C
Rosa26 wt
Dbh-iCRE
(DBH primer)
Dbh-iCRE

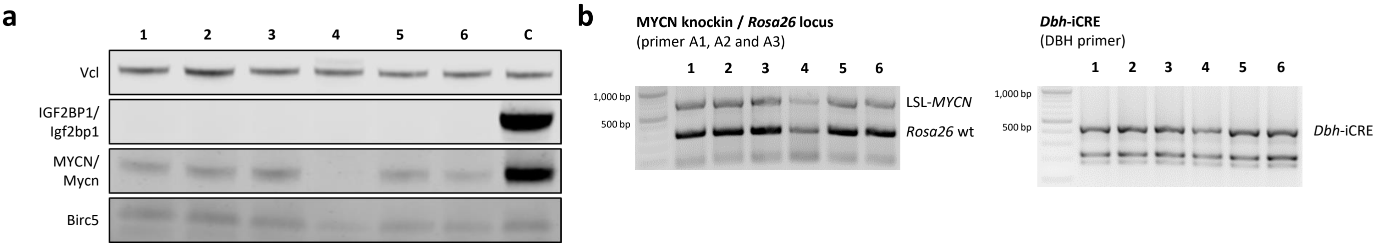
a
1 2 3 4 5 6 C
Vcl
IGF2BP1/
Igf2bp1
MYCN/
Mycn
Birc5
b
MYCN knockin / Rosa26 locus
(primer A1, A2 and A3)
1 2 3 4 5 6
1,000 bp
500 bp
LSL-MYCN
Rosa26 wt
Dbh-iCRE
(DBH primer)
1 2 3 4 5 6
1,000 bp
500 bp
Dbh-iCRE

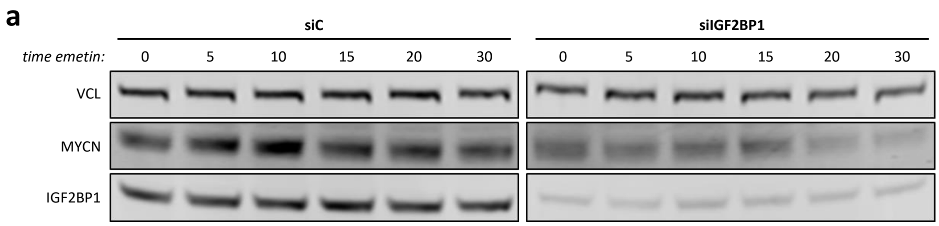
a
siC
siIGF2BP1
time emetin:
0
5
10
15
20
30
VCL
MYCN
IGF2BP1

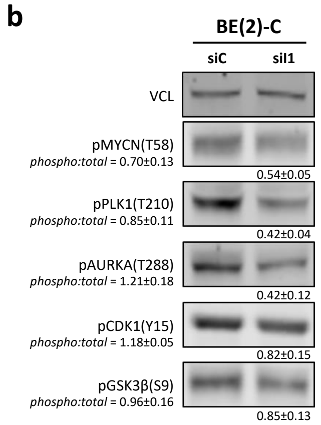
b
BE(2)-C
siC
siI1
VCL
pMYCN(T58)
phospho:total = 0.70±0.13
0.54±0.05
pPLK1(T210)
phospho:total = 0.85±0.11
0.42±0.04
pAURKA(T288)
phospho:total = 1.21±0.18
0.42±0.12
pCDK1(Y15)
phospho:total = 1.18±0.05
0.82±0.15
pGSK3β(S9)
phospho:total = 0.96±0.16
0.85±0.13

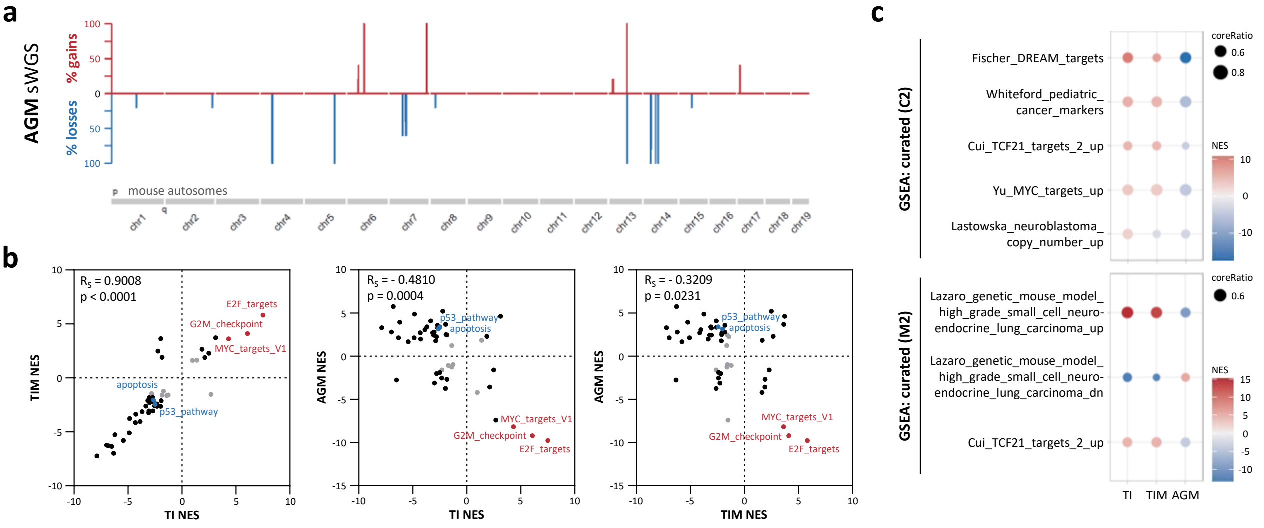
a
AGM sWGS
% gains
% losses
100
50
0
50
100
mouse autosomes
p
q
chr1
chr2
chr3
chr4
chr5
chr6
chr7
chr8
chr9
chr10
chr11
chr12
chr13
chr14
chr15
chr16
chr17
chr18
chr19
b
R_S = 0.9008
p < 0.0001
E2F_targets
G2M_checkpoint
MYC_targets_V1
apoptosis
p53_pathway
TIM NES
TI NES
R_S = - 0.4810
p = 0.0004
AGM NES
TI NES
R_S = - 0.3209
p = 0.0231
AGM NES
TIM NES
c
GSEA: curated (C2)
Fischer_DREAM_targets
Whiteford_pediatric_
cancer_markers
Cui_TCF21_targets_2_up
Yu_MYC_targets_up
Lastowska_neuroblastoma_
copy_number_up
coreRatio
0.6
0.8
NES
10
0
-10
GSEA: curated (M2)
Lazaro_genetic_mouse_model_
high_grade_small_cell_neuro-
endocrine_lung_carcinoma_up
Lazaro_genetic_mouse_model_
high_grade_small_cell_neuro-
endocrine_lung_carcinoma_dn
Cui_TCF21_targets_2_up
coreRatio
0.6
NES
15
10
5
0
-5
-10
TI
TIM
AGM
